# Learning from Drops: AI-Guided Integration of Liquid Biopsy Features in Cancer Studies

**DOI:** 10.64898/2026.05.12.724535

**Authors:** M. Andueza, P. Villoslada-Blanco, B. de Dreuille, L. Alonso, S. Sabroso-Lasa, K. Pantel, C. Alix-Panabières, E. López de Maturana, N. Malats

**Author notes:** **Correspondence to:** <u>Núria Malats</u>, Genetic and Molecular Epidemiology Group, Spanish National Cancer Research Centre (CNIO), C/Melchor Fernandez Almagro 3, 28029-Madrid, Spain.

## Abstract

Cancer is a major global health issue with rising incidence and mortality. Early detection, tumor characterization, and disease surveillance are crucial for timely and effective treatment, ultimately reducing mortality rates. Liquid biopsy (LB) has emerged as a valuable detection tool offering a non-invasive method to determine tumor-derived biomarkers in body fluids with demonstrated translational potential. To increase biomarker sensitivity, high-throughput sequencing platforms deliver massive volumes of data. Artificial Intelligence (AI) is pivotal in enabling huge and complex data integration. This contribution aims to assess the current state of integrative AI-based research in the LB field and provide methodological guidance. First, we conducted a PubMed search and found that the literature is sparse in studies integrating LB features, particularly by applying AI. When adopting the latter approach, defining the study objectives is crucial to guide the subsequent methodological aspects, including study design, patient selection criteria, sample size, nature of the LB features, and metadata to collect. Specifically, we propose strategies and tools for data preprocessing, including normalization and batch correction, as well as handling outliers and missing data. Furthermore, we recommend various Machine/Deep Learning approaches for feature selection techniques to ensure model robustness, and we highlight the importance of undergoing rigorous internal and external validations of the selected models. Assessing clinical utility and interpretability is often overlooked but fundamental for real-world implementation. In conclusion, we provide the LB scientific community with an AI-based methodological guidance to bridge the two fields and enhance the integrative analysis of LB features.

**Graphical abstract:** Workchart for multiomics integrative studies in the liquid biopsy field. **Note:** CTCs, circulating tumor cells; ctDNA, circulating tumor-DNA; TEPs, tumor-educated platelets; miRNA, microRNA; cfRNAs, cell-free RNAs.

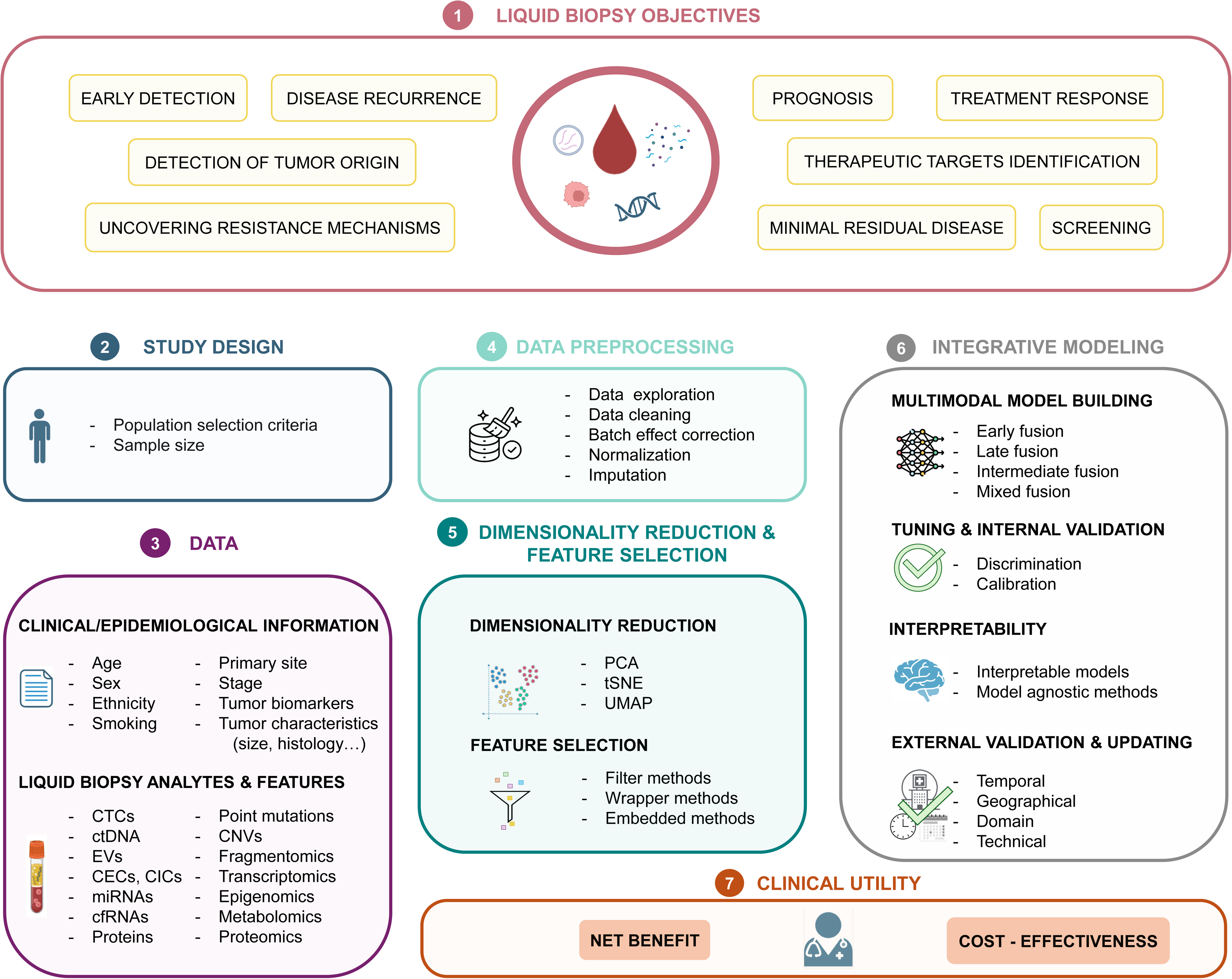

## Introduction

Cancer is a major health issue worldwide, being the second leading cause of death and showing a rapid increase in incidence and mortality ^1^. According to estimates from the World Health Organization in 2022, nearly 10 million people died from cancer that year, and there were an estimated 20 million new cancer cases ^2^. This number is predicted to reach 27 million cases by 2040 ^3^.

Early detection and prediction of tumor response through tumor profiling are crucial in cancer control, enabling both effective treatment of patients and a decrease in mortality rates ^4,5^. Indeed, in solid cancers, tumors are known to present a large heterogeneity, and various subtypes can be found within a cancer type. Moreover, the tumor can evolve both temporally and spatially with metastasis development, and new genomic alterations and epigenetic changes are acquired over tumor progression ^6,7^. Thus, molecular profiling of the tumor is essential to properly diagnose, treat, and monitor cancer patients in a personalized manner.

The traditional methods for characterizing the disease include imaging and tissue biopsies, the latter being used for both morphological and molecular profiling of the tumor. However, tissue biopsy is an invasive technique, and a single tumor biopsy may provide partial information due to intratumor heterogeneity, leading to the underassessment of the tumor’s genomics landscape ^7^. Considering these limitations and the increasing willingness to develop personalized medicine, new techniques for monitoring tumor evolution using biomarkers have been studied.

In particular, the concept of liquid biopsy (LB) was introduced in 2010 as a new diagnostic technique ^8^ and has generated great interest in recent years. This concept refers to the analysis of various biological fluids (blood, urine, saliva, and others), which contain circulating molecules and particles that can be shed by both primary tumor and metastasis, such as circulating tumor cells (CTCs), cell-free DNA (cfDNA), circulating tumor DNA (ctDNA), tumor-derived extracellular vesicles (EVs), tumor-educated platelets (TEPs), and microRNAs (miRNA), among others ^9,10^. Several studies have shown the potential of LB features analysis for early detection, tissue of origin (TOO) detection, minimal residual disease (MRD) monitoring, prognosis prediction, therapeutic targets identification, resistance mechanisms uncovering, and surveillance of treatment response ^11–18^. Using LB in clinical practice would allow less invasive and cost-efficient examinations for real-time monitoring of the patients^19–21^. However, the sensitivity of current LB assays is limited by several factors, including the low and variable levels of tumor DNA shed into the bloodstream, which differ between patients and fluctuate over time ^22,23^. Therefore, high-sensitivity and specificity models integrating multimodal LB features need to be built and properly evaluated, in addition to assessing their clinical utility.

In this regard, Artificial Intelligence (AI), including Machine Learning (ML) and Deep Learning (DL), can be highly helpful in handling the large amounts of complex data generated by the high-throughput biomarker platforms. Indeed, as single-candidate biomarker analysis is usually insufficient to predict a diagnosis or prognosis ^24–26^, ML and DL enable the simultaneous modeling of the effects of multiple biomarkers derived from different omics layers, accounting for their interactions. Consequently, integrating multiple biomarkers with AI can remarkably improve the performance of LB tests to predict various outcomes in cancer patients ^27,28^.

Here, we define multiomics integration as the process of combining molecular features originating either from different biological entities (e.g., ctDNA mutations and EV proteomics) or from different ‘omics’ layers within the same entity (e.g., ctDNA genomics and transcriptomics) into a single model. Although this integration typically involves tabular data, it aligns conceptually with multimodal ML, which focuses on combining diverse data types, such as images (radiomics, pathomics), clinical reports, and time series. From this perspective, multiomics integration can be seen as a specific instance of multimodal learning in the biomedical context ^29^. Multimodal integration builds on multimodal representation, translation, alignment, and co-learning, in addition to data fusion. Introducing these approaches to the field of health data analysis constitutes a challenge, but they will be highly valuable and lead to rapid advances in multiomics biomedical integration studies.

We documented the scarcity of integrative efforts for LB features in the scientific literature through a PubMed search of articles published from January 2020 to January 2026, as stated in the **Supplementary Methods**. This search, intended to identify representative studies rather than to constitute a systematic review, suggests that relatively few works have explored the integration of LB features across multiple molecular layers, probably because of the complexity of implementing AI approaches ^30^ (**Table S1**).

Although the search was limited to predefined terms (**see Supplementary Methods**) because of the narrative scope of the contribution, this effort enabled us to identify methodological gaps in applying AI-based integrative tools and in model evaluation within the LB field. Therefore, we aimed to provide methodological guidance for multilayer LB features integration through AI approaches. To this end, we covered aspects including study purpose and design, statistical characteristics of LB features, data preprocessing, dimensionality reduction and feature selection, multimodal integration with AI algorithms, and model evaluation for clinical practice. We exemplified each section with published data identified through the bibliographic search.

## Study purpose and design

As in any other study, the first step in designing a LB integration study is to define its primary **objective**. In this regard, LB biomarkers may benefit cancer patient management through early detection, TOO prediction, MRD monitoring, prognostication, therapeutic targets identification, resistance mechanisms uncovering, and treatment response surveillance^28^ Additionally, defining the necessary endpoints (primary and secondary) is crucial to ensure accurate and objective assessment. Poor selection of endpoints hinders the interpretation and implementation of the findings ^31^. When assessing prognosis, the time from diagnosis to death (i.e., overall survival) is considered the primary endpoint for oncologic clinical trials. Other endpoints currently considered in prognostic studies include disease-specific survival, progression-free survival (measured as the time from diagnosis until the first evidence of disease progression or death), and disease-free survival (measured as the time from diagnosis until the first evidence of disease recurrence or death) ^32^. If the study’s purpose is (early) diagnosis, the presence of the disease would be the expected outcome.

The research objectives, as well as the feasibility of conducting the study prospectively or retrospectively, determine its design. **Table S2** presents possible study designs related to each type of objective. Currently, many studies on LB are retrospective case-control studies, using already collected samples to measure LB biomarkers. This is the case for 18 out of the 25 studies identified in the literature search (**Table S1**). This type of design enables researchers to leverage previous efforts, making it less time-consuming and more cost-effective, albeit subject to potential bias. Moreover, epidemiological/clinical data may be incomplete or inadequate in answering the research questions appropriately. On the contrary, prospectively conducted studies would overcome those limitations by allowing real-time data collection under controlled conditions, thereby increasing the reliability and clinical relevance of LB biomarker validation.

Another key aspect of the study design relates to the definition of the **population**, including the **selection criteria** and the circuits needed to identify the subjects. Selection criteria are described in 22 out of the 25 identified studies. Moreover, estimating the number of individuals to be recruited (i.e., **sample size**) also impacts the model performance. Despite its importance, few studies mention this point ^33^. Only two papers from the bibliographic search estimated the sample size by conducting a power analysis after establishing a desired area under the curve (AUC) of 0.95 and 0.98 ^34,35^. A method proposed by Riley *et al*. ^36^ enables the calculation of sample size in clinical prediction models for binary, time-to-event, and continuous outcomes, utilizing a four-step procedure. Software for these calculations is available in the *psampsize package* in Stata or R. The package also estimates the minimum sample size required for external validation of clinical prediction models ^37,38^. However, these strategies primarily apply to classical models, where the number of parameters is limited. When integrating thousands of parameters by applying ML or DL, methodologies for sample size estimation are lacking. Therefore, in such scenarios, having more data typically leads to more robust results ^39^. Research is currently ongoing to determine the necessary sample size for these types of models. Recently, Goldenholz *et al*. ^40^ suggested applying a model-agnostic, open-source tool to evaluate sample size in ML clinical validation studies (Sample Size Analysis for Machine Learning, SSAML), which can be found at https://github.com/GoldenholzLab/SSAML.

Furthermore, the **information** to be collected in the study needs to be clearly defined. This encompasses a range of data, including demographic and epidemiological information, clinical characteristics of subjects, pathological or radiological details of the tumor, and patient follow-up. It is known that such traits can influence cancer detection, prognosis, and levels of biomarkers, and therefore their performance in outcome prediction. Specifically, age, gender, race, and ethnicity ^41–43^ significantly impact the prevalence, progression, and response to cancer detection technologies ^43–45^. Similarly, clinicopathological variables—including cancer type and stage, prior treatments, comorbidities, and genetic profile of both host and tumor—also affect the cancer prognosis ^35,46^. Nguyen *et al.* ^47^ also showed the influence of clinical features on model prediction by evaluating the potential confounding effect of gender, age, cancer burden, and cancer stage in their integrative model for multi-cancer early detection (MCED). Collecting all this information and considering these variables in the AI-based modeling as potential confounding factors, effect modifiers, or mediating variables certainly enhances the validity and accuracy of the model, making their results more robust, interpretable, clinically relevant, and generalizable to other populations ^44,45^. Another essential point in multicentric studies is the standardization of the workflow for information collection. The use of validated questionnaires ensures the reliability of the information gathered, thereby minimizing the use of different criteria in variable definition and categorization. Furthermore, studies have demonstrated that sampling conditions influence biomarker discovery in LB ^48^. Therefore, information regarding biosample collection, processing, and storage is necessary to ensure that these factors do not influence the results.

## Characteristics of liquid biopsy features

Liquid biopsy enables the exploration of numerous tumor-derived biomarkers present in body fluids. Among these, CTCs—tumor cells shed from the primary tumor and/or from distant metastases into the bloodstream—are well studied. Beyond cellular components, LB allows the study of other circulating biomarkers, including cell-free DNA and cell-free RNA. When derived from the tumor, these are referred to as circulating tumor DNA (ctDNA) and circulating tumor RNA (ctRNA), respectively. ctDNA consists of fragmented tumor-derived DNA found in the bloodstream, while ctRNA corresponds to RNA molecules released by tumor cells. Additionally, microRNAs (miRNAs)—small non-coding RNAs involved in gene regulation—and various circulating proteins, including those studied through secretomics (the analysis of secreted proteins), represent important analytes examined through LB. Other tumor-induced circulating cell types, such as circulating immune cells, circulating endothelial cells, cancer-associated fibroblasts, and TEPs, are also gaining increasing attention in oncological research and can be analyzed through LB. Extracellular vesicles, which carry molecular cargo such as proteins, RNA, and DNA, are another relevant class of biomarkers. Among them, exosomes ^49^—small EVs ranging from 30 to 150 nm and formed within the endosomal system—are widely studied. These circulating components have diverse cellular origins, which must be considered when addressing specific biological questions in cancer research. Moreover, they can be assessed through various analytical perspectives, including measuring their abundance or concentration, or by characterizing their molecular profiles using high-throughput technologies. Such analyses depend on both the origin and composition of each component. **Figure 1** illustrates these relationships, which are essential for determining variable types in multiomics integration studies. These distinctions also inform the statistical properties of the derived features and guide the selection of appropriate data processing steps. Further details are provided in **Table S3**.

**Figure 1.**
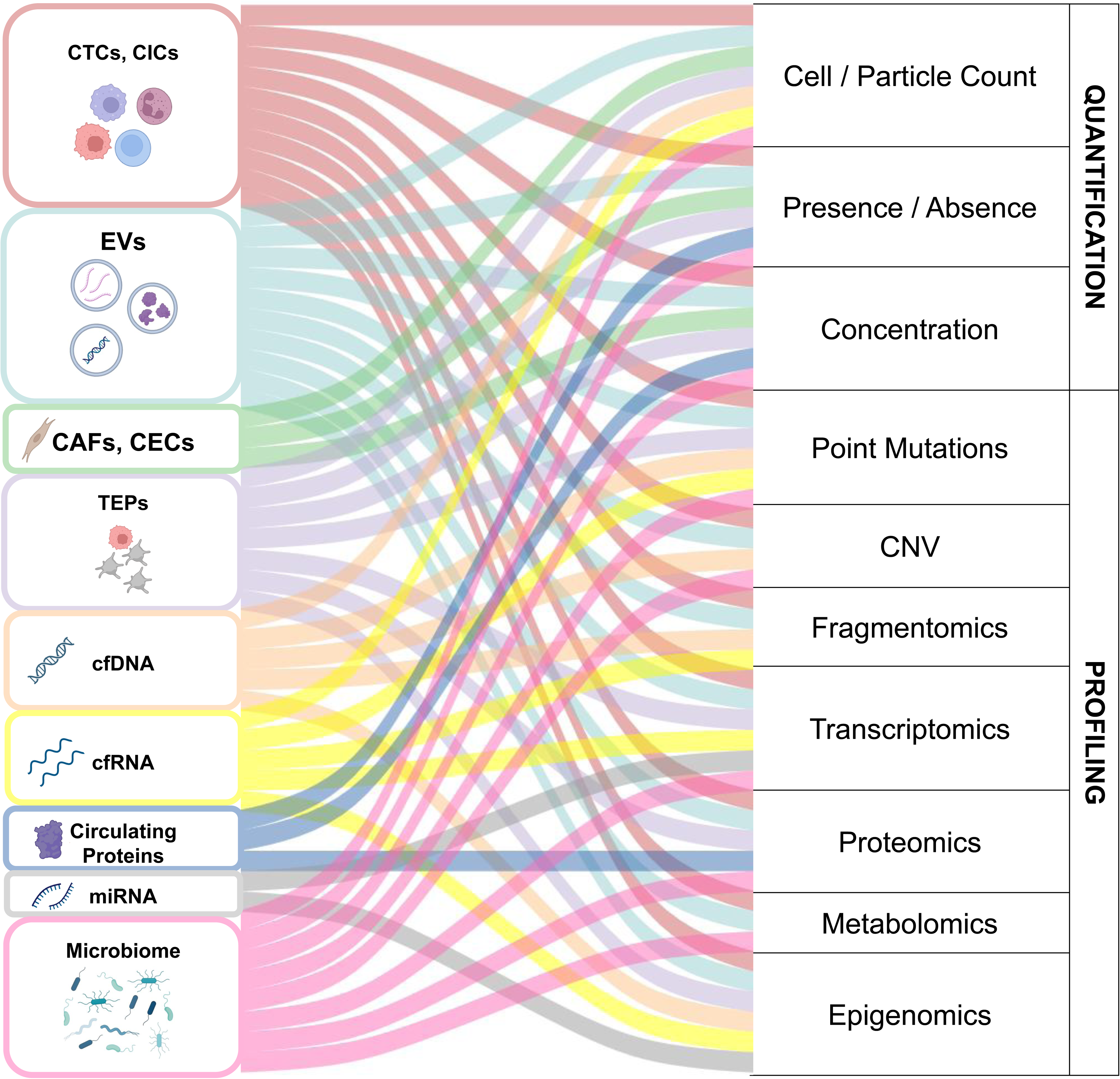
Correspondence between liquid biopsy components and omics layers/features. **Note:** CTCs: Circulating tumor cells, CICs: Circulating immune cells, EVs: Extracellular vesicles, CAFs: Cancer-associated fibroblasts, CECs: Circulating endothelial cells, cfDNA: circulating free DNA, miRNA: microRNA, CNV: Copy number variation.

## Data preprocessing

Understanding and visualizing data characteristics, also known as **data exploration**, is an initial step in data preprocessing and is particularly important for creating a robust dataset before modeling multimodal LB features using AI. This process is frequently followed by a **data cleaning** step, which involves detecting and correcting inconsistent records within the dataset. This step should be performed for each LB modality type and clinical/epidemiological variable. **Outlier handling** should also be considered at this point. The presence of **missing data** and incomplete records is another common challenge in multimodal integration. Imputation methods enable the completion of records to train AI models effectively. *MissForest* is a popular imputation method that enables the imputation of any type of data, including those with complex relationships between variables ^50^. It is a non-parametric method based on random forest (RF), which achieves high accuracy and can perform well with small datasets, but is highly computationally demanding. *K-nearest neighbors* (KNN) imputation models are also frequently used for numerical data imputation ^51^, and require no training. They perform well for small datasets whilst preserving the original data distribution. *Multiple Imputation by Chained Equations* (MICE) generates multiple imputed datasets to pool the results of the developed model, allowing for the incorporation of uncertainty associated with the actual values of the imputed variables ^52^. This method can be applied to various types of data, but it requires parameter tuning and becomes computationally intensive when handling many variables. Multiple imputation can be performed with the MICE package in R. Wang et al. ^53^ used MICE to impute data followed by sensitivity analysis. Although many studies perform imputation, very few assess the quality of the imputation itself or the impact on model performance of the distribution of missing data and its imputation.

Furthermore, zeros in the measures can be biological (truly absent from the data) or technical (introduced by the detection limit or library preparation steps) ^54^. Therefore, the lack of detection of a biomarker does not necessarily imply its absence in the sample. **Zero imputation** is used to accurately infer unseen values due to technical constraints, enabling for more reliable analysis and interpretation. There are different zero imputation methods, some developed for single-cell RNA sequencing data (*DrImpute, scImpute*), microarrays (*LLSimpute*), or compositional datasets (*zCompositions*) that vary in their methodologies ^55^ and are therefore more appropriate for specific LB data layers.

Even within the same data type, variability in biosample collection, processing, storage, or lab protocols—especially in retrospective studies—can introduce inconsistencies that compromise data reliability and comparability. Addressing potential batch effects is therefore essential to minimize technical variation in downstream analyses. **Table S4** describes various **batch effect correction** methods, along with a brief description of the input data type, their advantages, and disadvantages.

**Data normalization** is also a standard preprocessing step usually applied to standardize and scale the data to improve consistency. The nature of each layer must be considered when normalizing the data of an LB study that aims to integrate information from multiple features. As depicted in **Table S3**, most LB biomarkers are binary or continuous variables, with varying ranges. Methods such as Z-score standardization ^56^, max-min normalization, log-scaling, or decimal scaling have been proposed for normalizing LB features. ^54,55,57^

All the selected papers from our search considered more than one LB omics layer (analytes) containing multiple features. Hence, as high-dimensional data are computationally expensive to analyze, dimensionality reduction and feature selection must be performed at both single- and multiomics levels ^58–60^.

## Dimensionality reduction and feature selection

There are unsupervised and supervised methods for **dimensionality reduction**. In the case of single omics, *PCA (Principal Component Analysis), Independent Principal Component Analysis, UMAP (Uniform Manifold Approximation and Projection), or t-SNE (t-distributed Stochastic Neighbor Embedding)* constitute unsupervised methods, while *Linear Discriminant Analysis (LDA)* can be used as supervised dimensionality reduction. For multiomics data, unsupervised methods include *Canonical Correlation Analysis, MOFA (Multiomics Factor Analysis)*, or Joint and Individual Variation Explained, among others. Supervised methods include *Partial Least Squares, Multivariate Integration, or DIABLO (Data Integration Analysis for Biomarker discovery using Latent Variable approaches for Omics studies)*. The *mixOmics* ^61^ package in R offers many of the previously mentioned statistical methods for dimensionality reduction. Moreover, autoencoders or other DL methods are also suitable for this purpose.

**Feature selection** aims to identify the most informative and relevant biomarkers, improving model performance and interpretability. Feature selection techniques are categorized into filter, wrapper, and embedded methods. *Filter methods* evaluate feature importance based on intrinsic properties. They include Pearson’s correlation coefficient, F-statistic, Mutual Information, and Chi-squared statistic. *Wrapper methods* generate feature subsets using search algorithms. Strategies like recursive feature elimination, sequential selection algorithms, and meta-heuristic algorithms can be employed ^57,62^. For instance, the DEcancer framework ^63^ employed recursive feature elimination guided by RF variable importance to select a parsimonious multimodal biomarker panel that integrates protein biomarkers and ctDNA mutation features. This approach explicitly aimed to reduce redundancy while preserving predictive performance, highlighting the importance of systematic feature selection when combining heterogeneous liquid biopsy modalities. *Embedded methods* perform feature selection during model training process. These methods are specific to the adopted ML algorithm. Regularization methods like LASSO, Ridge, and Elastic Net (ENET) were used for feature selection in some selected papers ^47,57,64^. Tree-based methods, such as RF and gradient boosting, were also considered to rank the importance of various biomarkers ^47,64,65^. These methods are used to perform both feature selection and outcome prediction tasks simultaneously.

## Integrative modelling

This phase comprises four important and complex steps: 1. multimodal fusion, 2. model tuning and internal validation, 3. model interpretability, and 4. external validation and model updating. Below, we describe each step and provide lists of tools for the reader to consider when undertaking this endeavor.

### 1. Multimodal fusion

Also known as data fusion, must be implemented when building integration models, that combine information from multiple modalities (in this case, omics) to predict an outcome. Baltrusaitis *et al*. ^66^ classified the fusion techniques into model-based and model-agnostic approaches. The former models specifically address fusion in their construction and include *Multi-Kernel Learning*, *Graphical Models*, *Neural Networks*, or *Multi-View Learning* models, encompassing data fusion within these models. The latter performs data fusion independently of the model used. Moreover, approaches can be divided into early, late, and intermediate fusion. *Early fusion* (feature-level fusion) refers to the integration of all omics-derived features immediately after extraction, which can be achieved through concatenation, operations, or DL models. In *late fusion* (model-level fusion), models are built for each omics data layer and then integrated to produce a combined model output. More complex methodologies include *intermediate fusion* (also known as middle fusion or joint integration), where individual models first analyze the different layers before combining their outputs into a final prediction model. These terms are further represented in **Figure 2**. There are also mixed scenarios (*mixed fusion*) where approaches from the two strategies are combined ^67^.

**Figure 2.**
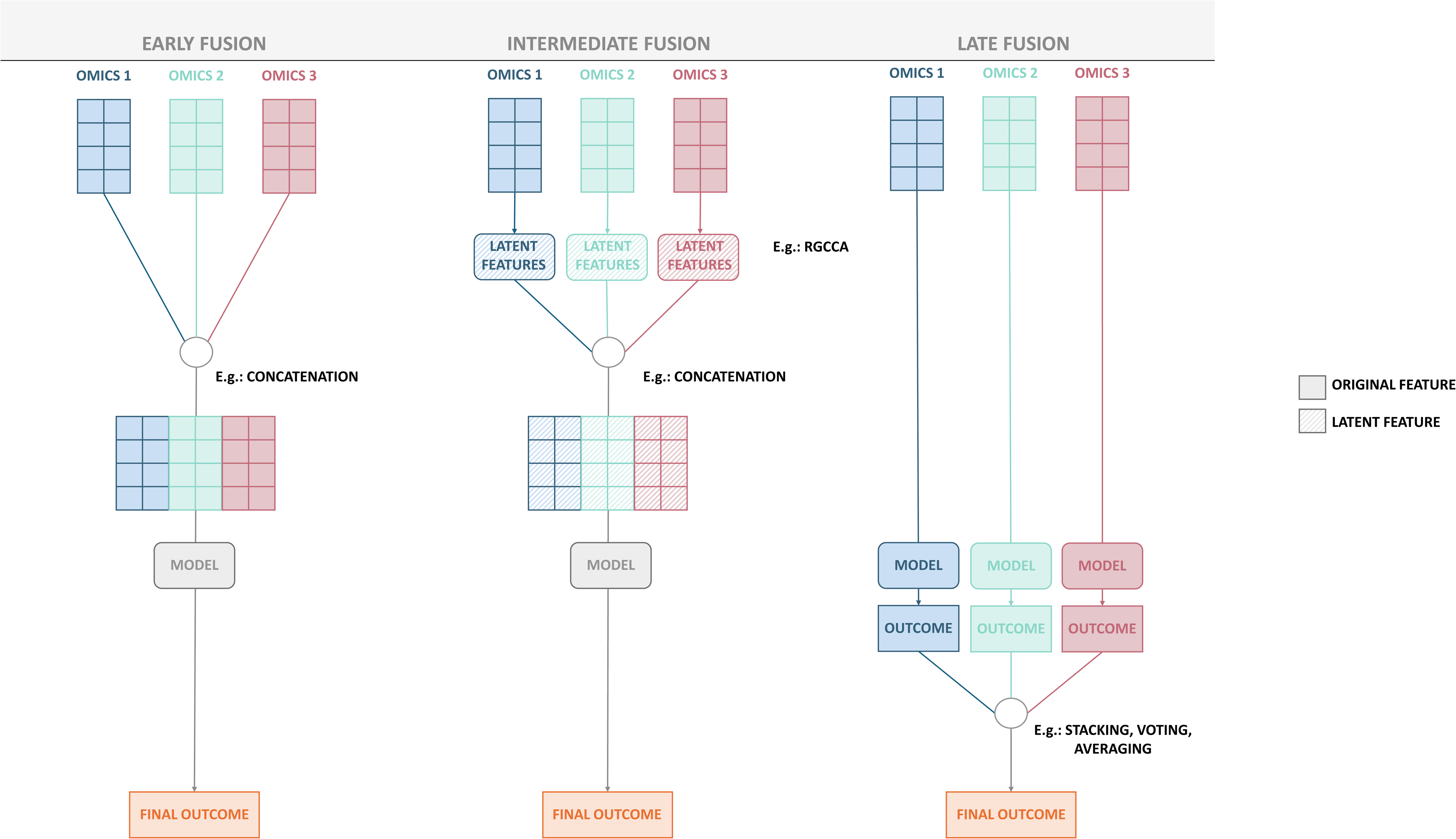
Data fusion approaches. **Note:** PCA: Principal component analysis.

To illustrate the aforementioned terms, we consider the methodological approaches adopted in Nguyen *et al* ^47^. Their approach for multi-cancer early detection followed two strategies: first, they generated a single model with the raw data of the LB feature modalities (targeted methylation, genome-wide methylation, ctDNA copy number alterations, fragment length, and end motifs) followed by an evaluation of the three different ML approaches on the complete dataset. This is an example of an early fusion strategy using a concatenation of the single features. Second, they built an ensemble stacking model combining the outcomes of nine base models (one per LB layer) with Linear Regressions (LR). This is considered late fusion. For the first case, *Extreme Gradient Boosting (XGBoost)* outperformed the stacking ensemble model ^47^. This highlights the need to study different data fusion approaches, which can be decisive for improving model performance together with the selected AI model.

Finally, it is important to consider which **ML/DL algorithm** performs the best. LR, especially with regularization, offers built-in feature selection and interpretability. Kernel-based methods address high dimensionality and have been used, for instance, to integrate proteomics and transcriptomics in predicting pancreatic cancer progression during chemotherapy ^57^. Decision trees and ensemble methods like *RF* and boosting algorithms (e.g., *XGBoost*, *AdaBoost*) are also widely applied for their balance of performance and interpretability. *Support Vector Machines* handle complex decision boundaries through kernels. In more complex settings, artificial neural networks can model nonlinear relationships but require larger datasets. Within this group, *Multi-Layer Perceptrons* have outperformed simpler models in certain multiomics applications, such as lung cancer classification ^68^.

Other DL approaches, such as convolutional neural networks, attention-based models, and recurrent neural networks or long short-term memory networks (for longitudinal data) are emerging as powerful tools for integrating diverse omics layers. Transformers can integrate heterogeneous signals into a unified representation, using attention layers to highlight which molecular features or modalities carry the most predictive value for tasks. In the LB context, transformer-based models have been applied directly to cfDNA fragments for representation learning and cancer prediction (e.g., DECIDIA^69^). Besides, in recent years, large language models (LLMs) have emerged as a breakthrough in AI, leading to widely known tools such as ChatGPT. These models can understand and generate human-like text by learning patterns from vast amounts of data. Moreover, foundation models are large-scale ML models that are trained on broad, general-purpose data (such as text, images, or code) and can be adapted to a wide range of downstream tasks. LLM-based foundation models such as LLaMA have already been fine-tuned in the LB setting to predict cancer from cfDNA ^70^. While they have not yet been applied to LB multiomics integration, these approaches hold considerable promise for future applications.

### 2. Model tuning and internal validation

**Model tuning** is essential for selecting the optimal hyperparameters and model architecture (in DL). This process typically involves splitting the dataset into training, validation, and test sets. The model is trained on the training set, while the validation set is used for feature selection, hyperparameter optimization, architecture selection (in deep learning) and model selection (when several strategies are performed) (**Figure 3**). The final model is then evaluated on an independent test set, a step often referred to as internal validation, which assesses the model’s generalizability within the available data and ensures reproducibility. When traditional data splitting is not feasible due to a limited sample size, resampling methods such as bootstrapping and *k*-fold cross-validation offer robust performance estimates by repeatedly training and validating the model across different data subsets. In small-*n*/high-*p* settings, which are common in liquid biopsy studies, nested cross-validation (an approach using an inner loop for model selection and an outer loop for performance estimation) may be required to avoid optimistic bias when feature selection or hyperparameter tuning is performed.

**Figure 3.**
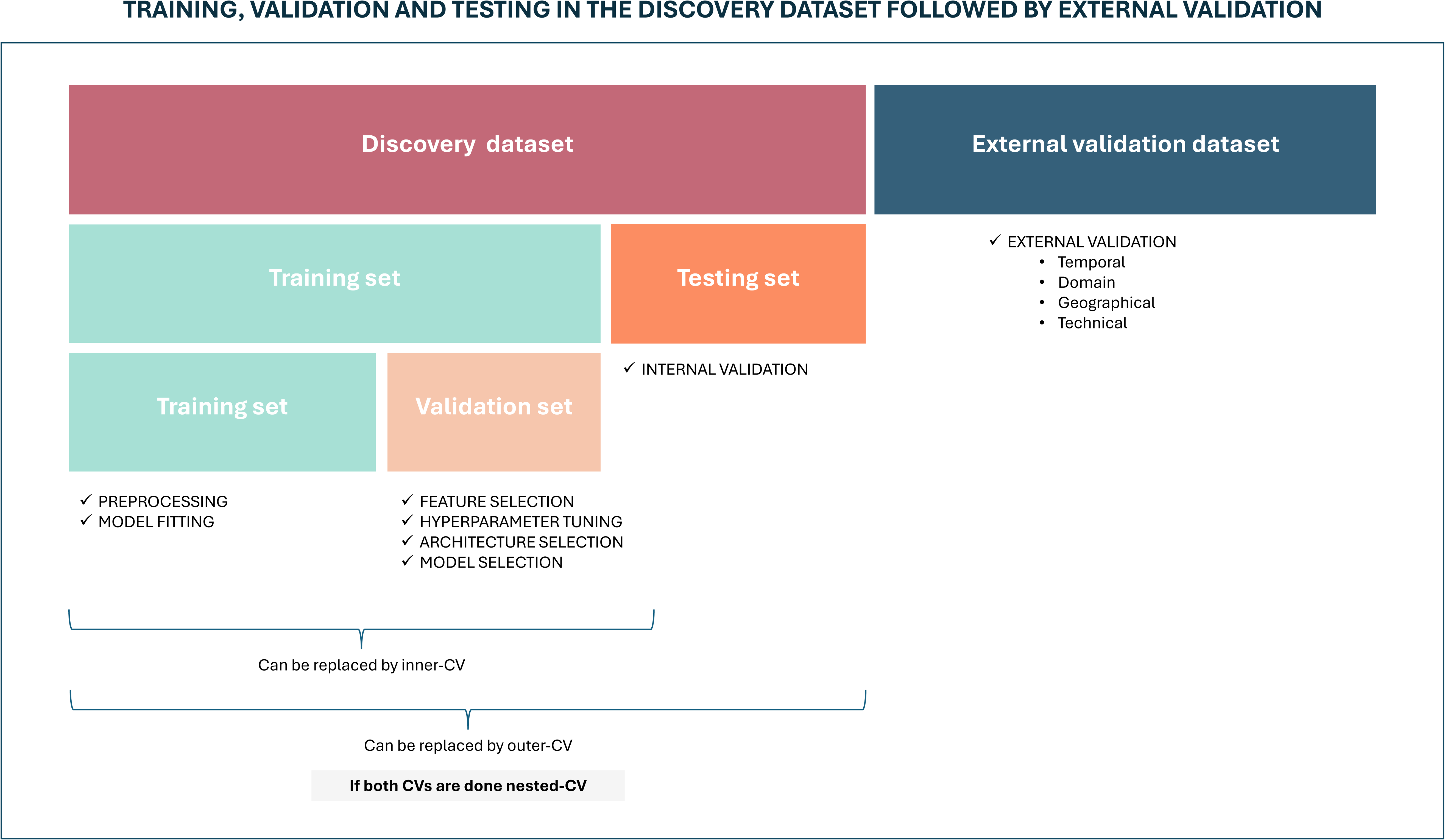
Schematic representation of data partitioning for AI model evaluation: includes training, validation, and testing in the discovery dataset, followed by external validation on an independent dataset.

When late-integration strategies are used, additional care is needed, as both the unimodal models (each with its own parameters) and the integration mechanism must be optimized. In such cases, model development should ensure that tuning of unimodal components and the final integration layer does not involve using data in the test set.

Two critical measures of the model validation are discrimination and calibration ^71^. These metrics provide unique insights into classification or regression performance, and the choice of metric depends on the specific application and the nature of the data. This ensures a comprehensive evaluation of models, leading to more accurate and reliable predictions. **Discrimination** measures the model’s ability to distinguish between different outcome states, such as the presence or absence of cancer. It evaluates how well the model distinguishes individuals with various outcomes based on their predicted risk scores. We have summarized different classification assessment methods in **Table S5**^72^, highlighting their methodologies, strengths, weaknesses, and applicability. One example of their application is proposed by Eledkawy *et al*. ^65^, who assessed the precise detection and classification of cancer using LB data and advanced ML techniques to analyze ctDNA mutations and protein biomarker concentrations. To evaluate the proposed model, they computed precision, recall, F1-score, accuracy, and AUC scores, showing the receiver operating characteristics (ROC) curve and confusion matrix results ^65^ and compared them with those obtained using previous methods on the same dataset.

On the other hand, **calibration** refers to the agreement between predicted probabilities and actual outcomes. A well-calibrated model provides accurate probability estimates across the entire range of predictions, ensuring that the predicted risk corresponds closely to the actual risk observed in the population ^73^. Calibration is crucial in clinical settings, where precise risk estimates can influence treatment decisions and patient management strategies. Standard metrics for assessing calibration include the observed-to-expected (O/E) ratio, calibration-in-the-large, calibration slope, and calibration plots. The O/E ratio compares the observed number of events to the expected number of events. Conversely, calibration-in-the-large evaluates the overall agreement between predicted probabilities and observed outcomes by adjusting the model intercept to correct for systematic over- or under-prediction (59,60). The calibration slope assesses whether the predicted probabilities are too extreme or not extreme enough ^73,74^. Finally, calibration plots compare predicted probabilities with observed outcomes, including the information of calibration-in-the-large and the calibration slope. These plots are useful for visually assessing how well the predicted probabilities align with actual outcomes across different risk levels ^73^.

Additionally, the Brier score provides a single measure of the accuracy of probabilistic predictions by computing the mean squared difference between predicted probabilities and actual outcomes, capturing both **calibration** and **discrimination** aspects.^75^

### 3. Model Interpretability

After selecting the best LB integrative model, evaluating its interpretability is crucial, especially in healthcare, where understanding decisions is key for biological interpretation and clinical implementation. Experts must fully understand the model’s pros and cons to translate them effectively into clinical practice.

For instance, consider building a model using proteomics and metabolomics data to predict disease recurrence. We could apply: 1) **interpretable models** offering transparency by design, as their decision rules or coefficients can be directly evaluated, being linear and logistic regressions, generalized linear models, generalized additive models, decision trees, and decision rules, the most interpretable ones; or 2) **more complex models,** some of which are seen as black-boxes, where interpretability methods must be implemented. After achieving high accuracy with the chosen model, it becomes crucial to understand which biomarkers are driving predictions. In the case of the first group of models, the interpretability is straightforward. For the most complex models, we could apply model-agnostic methods, which are flexible methods that work on any model and can be classified as global or local ^76^. **Global model-agnostic methods** provide insights into the overall logic of a model, while **local methods** explain the rationale behind specific individual predictions. For instance, SHAP (SHapley Additive exPlanations) is a model-agnostic local interpretation method that estimates the contribution of each feature to the output. This analysis may reveal that proteomic features in our example strongly influence predictions, highlighting key biomarkers and improving the model’s clinical relevance and cost-effectiveness. In fact, Wang et al.^53^ used SHAP values to rank model features, finding MRD, treatment modality and baseline ctDNA status as the top 3 contributors of their NN-based PRIME model (Progression Risk prediction using Interpretable ML on ctDNA-based genomic Mutations, MRD, and clinical-therapeutic features). A summary of the agnostic methods is presented in **Table S6**.

### 4. External validation and model updating

**External validation** is a cornerstone of developing robust integrative LB models. This process ensures the model generalizes effectively to new, independent data and related settings ^77^. This type of validation is critical for ensuring the model’s applicability in real-world clinical environments. It can take several forms and is implemented on an external validation dataset (**Figure 3**). *Temporal validation* assesses the model’s performance on data collected in a distinct time period from the training set, ensuring its consistency and robustness over time. *Geographic validatio*n uses data from multiple locations to test the model’s adaptability to diverse patient populations and healthcare practices. *Domain validation* involves applying the model across different clinical settings, such as transitioning from secondary to primary care environments, to evaluate its broader utility. Finally, *technical validation* ensures reproducibility across platforms and robustness under varying technical conditions. Six of the selected papers in this review validated or partially validated their results with distinct methodologies (**Table S1**), with an increasing tendency in recent years. Chabon *et al*. ^35^ externally validated their LungClip model, which discriminates early-stage lung cancer patients from risk-matched controls, in a prospectively collected independent cohort. Eledkawy *et al*. ^65^ partially validated their results using an external database, adapting their approach to the available features. Like internal validation, models are externally evaluated through discrimination and calibration metrics. Often overlooked due to the scarcity of validation data, this step is fundamental to model generalizability.

Beyond external validation, **model updating** is often needed. Cancer research continually evolves with advancements in medical knowledge, clinical practices, the discovery of new biomarkers, and the development of innovative treatments. Updates ensure the incorporation of new information, correct calibration drifts, and maintain generalizability across populations and settings. This should be applied to ensure models remain robust, adaptable, and applicable over time ^77^.

## Clinical utility and economic impact evaluation

Clinical utility is a critical aspect when evaluating LB integrative models. The goal of any predictive model in healthcare is not just to produce accurate predictions but to positively impact clinical outcomes and patient management. It measures how well a model aids in clinical decision-making, improves patient outcomes, and is cost-effective in real-world settings ^77^, justifying its use over existing methods.

Impact studies evaluate the model’s actual effect of the model on clinical practice. They can be designed as randomized controlled trials, before-and-after studies, or other comparative studies that measure changes in clinical outcomes, patient behaviors, and healthcare utilization following implementation of the predictive model ^78^. Moreover, the potential impact of the model can also be assessed with several metrics that, although often discussed, few publications have measured in their work. The *net benefit* quantifies clinical benefit, penalizing true positive classifications with false positive classifications ^79^, as shown in Equation 1:

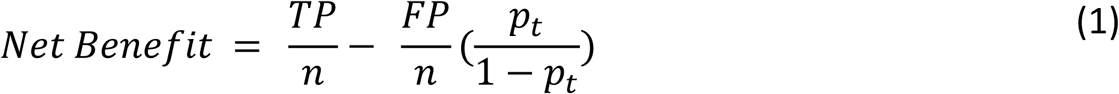

*Where TP:* True Positives*, FP:* False Positives*, n:* Total number of patients*, pt: threshold* where the expected benefit of treatment is equal to the expected benefit of avoiding treatment.

### Decision curve analysis

(DCA) evaluates the net benefit of a predictive model across different threshold probabilities (risk levels at which a clinician would choose to intervene), identifying when the model provides more benefit than harm compared to default strategies. The probability threshold is conceptually related to the minimum positive predictive value (PPV, the proportion of individuals with a positive test who truly have the disease) for a test to be clinically useful. PPV critically depends on disease prevalence in the screened population, in addition to test sensitivity and specificity. To illustrate these concepts, consider an AI-based LB model developed for early cancer detection, showing an AUC of 0.90 (high discrimination). A clinical deployment of this test would require selecting an operating point aligned with the intended use. If we want to use this AI model for population-level screening, a very high specificity threshold may be prioritized to minimize false positives and unnecessary downstream diagnostic procedures. Considering an operating point with 95% specificity and 60% sensitivity and a disease prevalence of 1% in the population to be screened, the AI model would identify approximately 6 true positives and 50 false positives per 1,000 screened individuals, which represents a substantial downstream diagnostic burden despite the high AUC of the AI model.

From a purely statistical perspective, one might select the operating point that maximizes the Youden index (sensitivity + specificity − 1), which identifies the threshold that optimizes overall discrimination. However, this approach does not account for disease prevalence, downstream consequences of false positives, or the clinical trade-offs inherent to screening decisions. Therefore, thresholds derived from the Youden index may not align with clinically meaningful risk thresholds, particularly in low-prevalence settings.

DCA at clinically plausible risk thresholds (e.g., 1–2%) can then reveal whether this trade-off provides a **net benefit** compared with “screen-all” or “screen-none” strategies. This example illustrates why AUC alone is insufficient and why explicit evaluation of operating points, calibration, and clinical utility metrics is essential for translational relevance. By integrating clinical consequences into the evaluation, DCA offers a nuanced understanding of the model impact on decision-making processes ^80^(79).

Besides, incorporating **economic impact evaluation**, such as cost-effectiveness analysis, helps to understand the financial implications of implementing the model in clinical practice. This involves comparing the costs and health outcomes of using the model against the standard of care, often using metrics like cost per quality-adjusted life year gained ^81^.

##### Box 1. Common pitfalls in AI-based integration of multi-layer LB data and recommended best practices

| Pitfall | Best practice |
| --- | --- |
| **Data leakage during data preprocessing**<br>Overly optimistic performance can arise when data preprocessing steps are conducted using the full dataset prior to data splitting | **Perform data preprocessing on the training data**<br>All steps should be learned on the training data and then applied unchanged to the test sets |
| **Improper model tuning**<br>Hyperparameter optimization performed outside the validation framework can lead to biased performance estimates, particularly in small-sample, high-dimensional settings typical of liquid biopsy studies | **Tune the models on the validation data**<br>Ensure that no data used for final evaluation influences model tuning decisions. The test set should remain completely independent and be used only for final performance evaluation |
| **Limited interpretability and reporting**<br>Reporting performance metrics without calibration assessment, uncertainty estimates, or a clear description of validation procedures limits clinical interpretability and transparency | **Ensure transparent reporting and interpretability**<br>Assess calibration, apply appropriate interpretability methods, and clearly document all preprocessing and validation steps |
| **Key principle:** All data-driven steps should be learned exclusively on the training data and applied **unchanged** to the test data* |  |

*\*In practice, certain preprocessing steps may be challenging to repeat within each CV fold due to computationally intensive procedures or methodological characteristics of particular omics data types. When such constraints arise, investigators should justify the chosen strategy and discuss the potential impact on performance estimates.*

## Challenges and future perspectives

This work provides the first methodological guidance for integrating AI-based models into the LB scenario and for addressing methodological gaps in cancer-related LB studies, including AI-integrationof multiomics/multimodal data. Data integration in the oncological field presents important challenges, including *(i)* securing high-quality and reliable data, which is essential for downstream analyses and for drawing biologically meaningful conclusions; *(ii)* adequate data preprocessing and exploratory analysis, often overlooked or vaguely described, which are critical steps to ensure data homogeneity and comparability across omics types, platforms, and studies; *(iii)* leveraging AI, implying the exploration of a variety of models, the evaluation of their performance using robust metrics, and the selection of the most appropriate approach based on the dataset characteristics and research goals; *(iv)* external validation and model calibration, often overlooked but essential to ensure generalizability and clinical applicability. Whereas internal evaluation is commonly performed, external validation using independent datasets, along with proper calibration and clinically meaningful metrics, is critical for translating models into real-world settings. *(v)* Model interpretability is crucial, as black-box models might be unsuitable for clinical applications. We highlight tools and strategies that enhance interpretability and understanding, particularly important when integrating multiple omics layers, where insights must be biologically and clinically meaningful. *(vi)* Models should be regularly updated to incorporate new information, correct calibration drifts, and maintain generalizability across populations and settings, ensuring long-term robustness, adaptability, and clinical relevance. *(vii)* Although not covered here, data privacy and ethical concerns are critical. The handling of multiomics data, particularly when linked to clinical information, raises concerns about patient privacy, data security, and informed consent. The fact that Ethical, Legal and Social Implications (ELSI) processes are not standardized and are highly fragmented (e.g., the European Union General Data Protection Regulation (GDPR) is differently interpreted by different member states) leads to significant delays in the project’s start, blocking the integration of data from larger research networks. Stringent harmonization of approval processes is, therefore, urgently needed. Aligning with the FAIR principles ^82^ is also essential for promoting transparency, responsible reuse, and scientific integrity while protecting participant confidentiality. Moreover, AI tools in healthcare should adhere to the FUTURE principles—Fair, Universal, Traceable, Usable, Robust, and Explainable ^83^. This guideline provides strategies to ensure these principles are upheld in LB integration studies.

In recent years, LLMs have demonstrated remarkable capabilities not only for data understanding and classification but also for generation. This has given rise to the field of generative AI (GenAIin which various models are used to generate new data across multiple modalities. Beyond natural language, GenAI now enables the creation of images, audio, and even complex biological or molecular data. In the context of multiomics LB research, such generative capabilities could allow for data augmentation, imputation of missing layers or features, synthetic data generation, and even foundation model building, opening new possibilities for liquid biopsy multiomics analysis. An early step in this direction is illustrated by Orion ^84^, a variational autoencoder applied to circulating non-coding RNAs, which leverages generative sampling in the latent space to augment training representations and improve cancer classification. This demonstrates how generative modeling concepts can enhance multiomics analysis even before full-scale GenAI is applied.

## Conclusion

While still an emerging field, an increasing number of studies are beginning to integrate data from multiple layers of LB omics to address critical questions in oncology. This review bridges LB and AI methodologies, offering guidance to ensure that studies are well-designed, that data are appropriately analyzed, and that conclusions are biologically relevant and clinically meaningful.

As a concise set of practical recommendations, researchers are encouraged to follow the steps outlined below:

1. **Clearly define the study objective and clinical context**, including the intended application (e.g., early detection, prognosis, treatment response) and the target population.
2. Understand the **biological and statistical properties of the data**, including feature types, distributions, sparsity, and potential sources of technical variability.
3. **Adopt an appropriate data-splitting and validation strategy**, choosing between train-test splits, cross-validation, or nested cross-validation depending on sample size and modeling complexity.
4. **Ensure that, when possible, preprocessing steps are performed on training data**, including outlier handling, missing value imputation, batch-effect correction, normalization, dimensionality reduction, and feature selection, and then applied to validation or test sets.
5. **Select an integration strategy and modeling approach aligned with the study objective**, considering early, intermediate, or late fusion and appropriate model classes (e.g., linear models, tree-based methods, DL).
6. **Report performance metrics that are appropriate for the task and data characteristics**, including discrimination and calibration, and interpret them in the context of class imbalance and clinical relevance.
7. **Incorporate model interpretability analyses**, using appropriate global and/or local explainability methods to assess feature contributions, support biological plausibility, and facilitate clinical trust.
8. **Whenever possible, perform external validation and model updating, and assess clinical utility**, as these steps are critical for evaluating generalizability.

We believe that adherence to these principles will improve reproducibility, robustness, and translational relevance of AI-based integrative LB studies.

## Acknowledgements

Not applicable.

## Author Contributions

**M. Andueza:** Conceptualization; literature search; literature synthesis; figure creation; writing – original draft preparation; writing – review and editing.

**P. Villoslada-Blanco:** Conceptualization; literature search; literature synthesis; writing – original draft preparation; writing – review and editing.

**B. de Dreuille:** Conceptualization; literature search; literature synthesis; writing – original draft preparation; writing – review and editing.

**L. Alonso:** Conceptualization; writing – original draft preparation; writing – review and editing.

**S. Sabroso-Lasa:** Conceptualization; writing – original draft preparation; writing – review and editing.

**K. Pantel:** Writing – review.

**C. Alix-Panabières:** Writing – review.

**E. López de Maturana:** Conceptualization; writing – original draft preparation; writing – review and editing.

**N. Malats:** Conceptualization; supervision; writing – review and editing

All authors approved the final version of the manuscript.

## Ethics Approval and Consent to Participate

Not applicable.

## Consent for Publication

Not applicable.

## Data and code Availability

Not applicable.

## Abbreviations used in this paper

AI: Artificial Intelligence
CCA: canonical correlation analysis
cfDNA: cell-free DNA
cfRNA: cell-free RNA
CTC: circulating tumor cell
ctDNA: circulating tumor DNA
ctRNA: circulating tumor RNA
DCA: decision curve analysis
DL: Deep Learning
EVs: extracellular vesicles
KNN: k-nearest neighbors
LASSO: least absolute shrinkage and selection operator
LB: liquid biopsy
LDA: linear discriminant analysis
LLM: large language model
LR: linear regression
MICE: multiple imputation by chained equations
miRNA: microRNA
ML: Machine Learning
MRD: minimal residual disease
NB: net benefit
O/E ratio: observed-to-expected ratio
RF: random forest
TEP: tumor-educated platelet
TOO: tissue of origin
XGBoost: extreme gradient boosting

## References

1. Sung H, Ferlay J, Siegel RL, et al. Global cancer statistics 2020: GLOBOCAN estimates of incidence and mortality worldwide for 36 cancers in 185 countries. CA Cancer J Clin. 2021;71(3). doi:10.3322/caac.21660

2. World Health Organization. Global cancer burden growing, amidst mounting need for services. Accessed July 3, 2024. https://www.who.int/news/item/01-02-2024-global-cancer-burden-growing--amidst-mounting-need-for-services

3. Wild, C. P.; Weiderpass, E.; Stewart BW. World Cancer Report: Cancer Research for Cancer Prevention. Vol 199. 2020.

4. World Health Organization. Guide to Early Cancer Diagnosis.

5. Loud JT, Murphy J. Cancer Screening and Early Detection in the 21st Century. Semin Oncol Nurs. W.B. Saunders. 2017;33(2):121–128. doi:10.1016/j.soncn.2017.02.002

6. Dumbrava EI, Meric-Bernstam F. Personalized cancer therapy-leveraging a knowledge base for clinical decision-making. Cold Spring Harb Mol Case Stud. Cold Spring Harbor Laboratory Press. 2018;4(2). doi:10.1101/mcs.a001578

7. Gerlinger M, Rowan AJ, Horswell S, et al. Intratumor heterogeneity and branched evolution revealed by multiregion sequencing. N Engl J Med. 2012;366(10):883–892. doi:10.1056/nejmoa1113205

8. Pantel K, Alix-Panabières C. Circulating tumour cells in cancer patients: Challenges and perspectives. Trends Mol Med. 2010;16(9). doi:10.1016/j.molmed.2010.07.001

9. Alix-Panabières C, Pantel K. Liquid biopsy: From discovery to clinical application. Cancer Discov. 2021;11(4):858–873. doi:10.1158/2159-8290.CD-20-1311

10. Lone SN, Nisar S, Masoodi T, et al. Liquid biopsy: a step closer to transform diagnosis, prognosis and future of cancer treatments. Mol Cancer. 2022;21(1). doi:10.1186/s12943-022-01543-7

11. Liu MC, Oxnard GR, Klein EA, et al. Sensitive and specific multi-cancer detection and localization using methylation signatures in cell-free DNA. Annals of Oncology. 2020;31(6). doi:10.1016/j.annonc.2020.02.011

12. Chaudhuri AA, Chabon JJ, Lovejoy AF, et al. Early detection of molecular residual disease in localized lung cancer by circulating tumor DNA profiling. Cancer Discov. 2017;7(12). doi:10.1158/2159-8290.CD-17-0716

13. Kilgour E, Rothwell DG, Brady G, Dive C. Liquid Biopsy-Based Biomarkers of Treatment Response and Resistance. Cancer Cell. 2020;37(4). doi:10.1016/j.ccell.2020.03.012

14. De Rubis G, Rajeev Krishnan S, Bebawy M. Liquid Biopsies in Cancer Diagnosis, Monitoring, and Prognosis. Trends Pharmacol Sci. 2019;40(3). doi:10.1016/j.tips.2019.01.006

15. Pantel K, Alix-Panabières C. Minimal residual disease as a target for liquid biopsy in patients with solid tumours. Nat Rev Clin Oncol. 2024;22(1). doi:10.1038/S41571-024-00967-Y,

16. Alix-Panabières C, Pantel K. Advances in liquid biopsy: From exploration to practical application. Cancer Cell. 2025;43(2):161–165. doi:10.1016/j.ccell.2024.11.009

17. Lennon AM, Buchanan AH, Kinde I, et al. Feasibility of blood testing combined with PET-CT to screen for cancer and guide intervention. Science (1979). 2020;369. doi:10.1126/science.abb9601

18. Reinert T, Henriksen TV, Christensen E, et al. Analysis of plasma cell-free DNA by ultradeep sequencing in patients with stages i to III colorectal cancer. JAMA Oncol. 2019;5(8). doi:10.1001/jamaoncol.2019.0528

19. Englmeier F, Bleckmann A, Brückl W, Griesinger F, Fleitz A, Nagels K. Clinical benefit and cost-effectiveness analysis of liquid biopsy application in patients with advanced non-small cell lung cancer (NSCLC): a modelling approach. J Cancer Res Clin Oncol. 2023;149(4):1495–1511. doi:10.1007/s00432-022-04034-w

20. Kramer A, Greuter MJE, Schraa SJ, et al. Early evaluation of the effectiveness and cost-effectiveness of ctDNA-guided selection for adjuvant chemotherapy in stage II colon cancer. Ther Adv Med Oncol. 2024;16. doi:10.1177/17588359241266164

21. Liu S, Graves N, Tan AC. The cost-effectiveness of including liquid biopsy into molecular profiling strategies for newly diagnosed advanced non-squamous non-small cell lung cancer in an Asian population. Lung Cancer. 2024;191. doi:10.1016/j.lungcan.2024.107794

22. Connal S, Cameron JM, Sala A, et al. Liquid biopsies: the future of cancer early detection. J Transl Med. 2023;21(1). doi:10.1186/s12967-023-03960-8

23. Menna G, Piaser Guerrato G, Bilgin L, Ceccarelli GM, Olivi A, Della Pepa GM. Is There a Role for Machine Learning in Liquid Biopsy for Brain Tumors? A Systematic Review. Int J Mol Sci. 2023;24(11). doi:10.3390/ijms24119723

24. Esposito A, Criscitiello C, Locatelli M, Milano M, Curigliano G. Liquid biopsies for solid tumors: Understanding tumor heterogeneity and real time monitoring of early resistance to targeted therapies. Pharmacol Ther. 2016;157. doi:10.1016/j.pharmthera.2015.11.007

25. Burrell RA, McGranahan N, Bartek J, Swanton C. The causes and consequences of genetic heterogeneity in cancer evolution. Nature. 2013;501(7467). doi:10.1038/nature12625

26. Fu Q, Schoenhoff FS, Savage WJ, Zhang P, van Eyk JE. Multiplex assays for biomarker research and clinical application: Translational science coming of age. Proteomics Clin Appl. 2010;4(3). doi:10.1002/prca.200900217

27. Ko J, Baldassano SN, Loh PL, Kording K, Litt B, Issadore D. Machine learning to detect signatures of disease in liquid biopsies-a user’s guide. Lab Chip. 2018;18(3). doi:10.1039/c7lc00955k

28. Chen G, Zhang J, Fu Q, Taly V, Tan F. Integrative analysis of multi-omics data for liquid biopsy. Br J Cancer. 2023;128(4):505–518. doi:10.1038/s41416-022-02048-2

29. Briscik M, Tazza G, Dillies MA, Vidács L, Dejean S. Supervised multiple kernel learning approaches for multi-omics data integration. BioData Min. 2024;17(1). doi:10.1186/S13040-024-00406-9

30. Liu L, Chen X, Petinrin OO, et al. Machine learning protocols in early cancer detection based on liquid biopsy: A survey. 2021;11(7). doi:10.3390/life11070638

31. McLeod C, Norman R, Litton E, Saville BR, Webb S, Snelling TL. Choosing primary endpoints for clinical trials of health care interventions. Contemp Clin Trials Commun. 2019;16:100486. doi:10.1016/j.conctc.2019.100486

32. Delgado A, Guddati AK. Clinical endpoints in oncology - a primer. Am J Cancer Res. 2021;11(4):1121.

33. Dhiman P, Ma J, Qi C, et al. Sample size requirements are not being considered in studies developing prediction models for binary outcomes: a systematic review. BMC Med Res Methodol. 2023;23(1):1–11. doi:10.1186/s12874-023-02008-1

34. Wang L, Zhang M, Pan X, et al. Integrative serum metabolic fingerprints based multi-modal platforms for lung adenocarcinoma early detection and pulmonary nodule classification. Adv Sci (Weinh). 2022;9(34):2203786. doi:10.1002/advs.202203786

35. Chabon JJ, Hamilton EG, Kurtz DM, et al. Integrating genomic features for non-invasive early lung cancer detection. Nature. 2020;580(7802):245–251. doi:10.1038/s41586-020-2140-0

36. Riley RD, Ensor J, Snell KIE, et al. Calculating the sample size required for developing a clinical prediction model. BMJ. 2020;368. doi:10.1136/bmj.m441

37. Riley RD, Debray TPA, Collins GS, et al. Minimum sample size for external validation of a clinical prediction model with a binary outcome. Stat Med. 2021;40(19):4230–4251. doi:10.1002/sim.9025

38. Riley RD, Collins GS, Ensor J, et al. Minimum sample size calculations for external validation of a clinical prediction model with a time-to-event outcome. Stat Med. 2022;41(7):1280–1295. doi:10.1002/sim.9275

39. Tsegaye B, Snell KIE, Archer L, et al. Larger sample sizes are needed when developing a clinical prediction model using machine learning in oncology: methodological systematic review. doi:10.1016/j.jclinepi.2025.111675

40. Goldenholz DM, Sun H, Ganglberger W, Westover MB. Sample size analysis for Machine Learning clinical validation studies. Biomedicines. 2023;11(3). doi:10.3390/biomedicines11030685

41. Bray F, Laversanne M, Sung H, et al. Global cancer statistics 2022: GLOBOCAN estimates of incidence and mortality worldwide for 36 cancers in 185 countries. CA Cancer J Clin. 2024;74(3):229–263. doi:10.3322/CAAC.21834

42. Boffetta P, Nyberg F. Contribution of environmental factors to cancer risk. Br Med Bull. 2003;68:71–94. doi:10.1093/bmp/ldg023

43. Colditz GA, Wei EK. Preventability of cancer: the relative contributions of biologic and social and physical environmental determinants of cancer mortality. Annu Rev Public Health. 2012;33:137–156. doi:10.1146/annurev-publhealth-031811-124627

44. Molina-Montes E, Coscia C, Gómez-Rubio P, et al. Deciphering the complex interplay between pancreatic cancer, diabetes mellitus subtypes and obesity/BMI through causal inference and mediation analyses. Gut. 2021;70(2):319–329. doi:10.1136/gutjnl-2019-319990

45. Molina-Montes E, Van Hoogstraten L, Gomez-Rubio P, et al. Pancreatic cancer risk in relation to lifetime smoking patterns, tobacco type, and dose–response relationships. Cancer Epidemiol Biomarkers Prev. 2020;29(5):1009–1018. doi:10.1158/1055-9965.EPI-19-1027

46. Weiss NS, Koepsell TD. Epidemiologic Methods: Studying the Occurrence of Illness. Oxford University Press; 2014. doi:10.1093/med/9780195314465.001.0001

47. Nguyen VTC, Nguyen TH, Doan NNT, et al. Multimodal analysis of methylomics and fragmentomics in plasma cell-free DNA for multi-cancer early detection and localization. 2023;12. doi:10.7554/elife.89083

48. Batool SM, Hsia T, Beecroft A, et al. Extrinsic and intrinsic preanalytical variables affecting liquid biopsy in cancer. Cell Rep Med. 2023;4(10):101196. doi:10.1016/j.xcrm.2023.101196

49. Van Niel G, D’Angelo G, Raposo G. Shedding light on the cell biology of extracellular vesicles. Nat Rev Mol Cell Biol. 2018;19(4). doi:10.1038/nrm.2017.125

50. Stekhoven DJ, Bühlmann P. MissForest--non-parametric missing value imputation for mixed-type data. Bioinformatics. 2012;28(1):112–118. doi:10.1093/bioinformatics/btr597

51. Troyanskaya O, Cantor M, Sherlock G, et al. Missing value estimation methods for DNA microarrays. Bioinformatics. 2001;17(6):520–525. doi:10.1093/bioinformatics/17.6.520

52. Austin PC, White IR, Lee DS, van Buuren S. Missing data in clinical research: A tutorial on multiple imputation. Can J Cardiol. 2021;37(9):1322–1331. doi:10.1016/j.cjca.2020.11.010

53. Wang Y, Xiang YB, Chen XW, et al. PRIME: an interpretable artificial intelligence model based on liquid biopsy improves prediction of progression risk in non-small cell lung cancer. Mil Med Res. 2025;12(1). doi:10.1186/s40779-025-00679-z

54. Silverman JD, Roche K, Mukherjee S, David LA. Naught all zeros in sequence count data are the same. Comput Struct Biotechnol J. 2020;18:2789–2798. doi:10.1016/j.csbj.2020.09.014

55. Baruzzo G, Patuzzi I, Di Camillo B. Beware to ignore the rare: how imputing zero-values can improve the quality of 16S rRNA gene studies results. BMC Bioinformatics. 2021;22(15):1–33. doi:10.1186/s12859-022-04587-0

56. Nakamura K, Zhu Z, Roy S, et al. An exosome-based transcriptomic signature for noninvasive, early detection of patients with pancreatic ductal adenocarcinoma: A multicenter cohort study. Gastroenterology. 2022;163(5):1252–1266.e2. doi:10.1053/j.gastro.2022.06.090

57. van Eijck CWF, Sabroso-Lasa S, Strijk GJ, et al. A liquid biomarker signature of inflammatory proteins accurately predicts early pancreatic cancer progression during FOLFIRINOX chemotherapy. Neoplasia. 2024;49. doi:10.1016/j.neo.2024.100975

58. Demler O V., Pencina MJ, D’Agostino RB. Impact of correlation on predictive ability of biomarkers. Stat Med. 2013;32(24):4196–4210. doi:10.1002/sim.5824

59. Bansal A, SullivanPepe M. When does combining markers improve classification performance and what are implications for practice? Stat Med. 2013;32(11):1877–1892. doi:10.1002/sim.5736

60. Pinsky PF, Zhu CS. Building multi-marker algorithms for disease prediction-the role of correlations among markers. Biomark Insights. 2011;6:83–93. doi:10.4137/bmi.S7513

61. Rohart F, Gautier B, Singh A, Lê Cao KA. mixOmics: An R package for ‘omics feature selection and multiple data integration. PLoS Comput Biol. 2017;13(11):e1005752. doi:10.1371/journal.pcbi.1005752

62. Genco E, Modena F, Sarcina L, et al. A single-molecule bioelectronic portable array for early diagnosis of pancreatic cancer precursors. Adv Mater. 2023;35(42):2304102. doi:10.1002/adma.202304102

63. Halner A, Hankey L, Liang Z, et al. DEcancer: Machine learning framework tailored to liquid biopsy based cancer detection and biomarker signature selection. iScience. 2023;26(5). doi:10.1016/j.isci.2023.106610

64. Pham TMQ, Phan TH, Jasmine TX, et al. Multimodal analysis of genome-wide methylation, copy number aberrations, and end motif signatures enhances detection of early-stage breast cancer. Front Oncol. 2023;13. doi:10.3389/fonc.2023.1127086

65. Eledkawy A, Hamza T, El-Metwally S. Precision cancer classification using liquid biopsy and advanced machine learning techniques. Sci Rep. 2024;14(1). doi:10.1038/s41598-024-56419-1

66. Baltrusaitis T, Ahuja C, Morency LP. Multimodal Machine Learning. IEEE Trans Pattern Anal Mach Intell. 2019;41(2):423–443. doi:10.1109/TPAMI.2018.2798607

67. Krones F, Marikkar U, Parsons G, Szmul A, Mahdi A. Review of multimodal machine learning approaches in healthcare. Inf Fusion. 2025;114:102690. doi:10.1016/j.inffus.2024.102690

68. Kwon HJ, Park UH, Goh CJ, et al. Enhancing lung cancer classification through integration of liquid biopsy multi-omics data with machine learning techniques. Cancers (Basel). 2023;15(18). doi:10.3390/cancers15184556

69. Liu J, Shen H, Yang Y, et al. Transformer-based representation learning and multiple-instance learning for cancer diagnosis exclusively from raw sequencing fragments of bisulfite-treated plasma cell-free DNA. Mol Oncol. 2024;18(11):2755. doi:10.1002/1878-0261.13745

70. Liu J, Shen H, Chen K, Li X. Large language model produces high accurate diagnosis of cancer from end-motif profiles of cell-free DNA. Brief Bioinform. 2024;25(5). doi:10.1093/bib/bbae430

71. Steyerberg EW. Clinical Prediction Models. Springer International Publishing; 2019. doi:10.1007/978-3-030-16399-0

72. Tharwat A. Classification assessment methods. Appl Comput Inform. 2018;17(1):168–192. doi:10.1016/j.aci.2018.08.003

73. Steyerberg EW, Vickers AJ, Cook NR, et al. Assessing the performance of prediction models: a framework for some traditional and novel measures. Epidemiology. 2010;21(1):128. doi:10.1097/ede.0b013e3181c30fb2

74. Van Calster B, McLernon DJ, Van Smeden M, et al. Calibration: The Achilles heel of predictive analytics. BMC Med. 2019;17(1):1–7. doi:10.1186/s12916-019-1466-7

75. Brier GW. Verification forecasts expressed in terms of probability. Mon Weather Rev. 1950;78(1). doi:10.1175/1520-0493(1950)078<0001:VOFEIT>2.0.CO;2

76. Interpretable Machine Learning. Accessed May 26, 2025. https://christophm.github.io/interpretable-ml-book/

77. Binuya MAE, Engelhardt EG, Schats W, Schmidt MK, Steyerberg EW. Methodological guidance for the evaluation and updating of clinical prediction models: a systematic review. BMC Medical Research Methodology 2022 22:1. 2022;22(1):1–14. doi:10.1186/s12874-022-01801-8

78. Collins GS, Reitsma JB, Altman DG, Moons KGM. Transparent reporting of a multivariable prediction model for individual prognosis or diagnosis (TRIPOD): The TRIPOD Statement. BMC Med. 2015;13(1):1–10. doi:10.1186/s12916-014-0241-z

79. Vickers AJ, Van Calster B, Steyerberg EW. Net benefit approaches to the evaluation of prediction models, molecular markers, and diagnostic tests. BMJ. 2016;352. doi:10.1136/bmj.I6

80. Vickers AJ, Elkin EB. Decision curve analysis: a novel method for evaluating prediction models. Med Decis Making. 2006;26(6):565–574. doi:10.1177/0272989X06295361

81. Moons KGM, Kengne AP, Grobbee DE, et al. Risk prediction models: II. External validation, model updating, and impact assessment. Heart. 2012;98(9):691–698. doi:10.1136/HEARTJNL-2011-301247

82. Wilkinson MD, Dumontier M, Aalbersberg IjJ, et al. The FAIR Guiding Principles for scientific data management and stewardship. Scientific Data 2016 3:1. 2016;3(1):1–9. doi:10.1038/sdata.2016.18

83. Lekadir K, Feragen A, Fofanah AJ, et al. FUTURE-AI: international consensus guideline for trustworthy and deployable artificial intelligence in healthcare. BMJ. 2025;388. doi:10.1136/BMJ-2024-081554

84. Karimzadeh M, Momen-Roknabadi A, Cavazos TB, et al. Deep generative AI models analyzing circulating orphan non-coding RNAs enable detection of early-stage lung cancer. Nat Commun. 2024;15(1):1–12. doi:10.1038/S41467-024-53851-9

