## Supplementary material for "Learning from Drops: AI-Guided Integration of Liquid Biopsy Features in Cancer Studies": Suplementary Material

Andueza et al. **Supplementary Material**

### Supplementary Methods

**Table S1.** Characteristics of the articles identified in the literature search.

**Table S2.** Correspondence between objectives and study designs.

**Table S3.** Liquid biopsy features characteristics.

**Table S4.** Summary of batch effect correction methods: characteristics, advantages, and limitations for application in liquid biopsy.

**Table S5**. Summary of model selection metrics: purpose, equation, advantages, and disadvantages.

**Table S6.** Summary of model-agnostic methods to interpret Machine Learning models.

**Supplementary Methods**

Details on the bibliographic search for this review are the following: the research was conducted via PubMed on research articles published with the terms ("cancer" OR "carcinoma" OR "tumor" OR "neoplasm") AND "liquid biopsy" AND ("machine learning" OR "deep learning" OR "artificial intelligence") AND ("early detection" OR "minimal residual disease" OR "prognosis"), excluding meta-analysis and reviews, and focusing on human studies. The search covered the period from January 2020 to January 2026 and provided 144 results. Of the 144 papers, 123 were discarded because the authors did not integrate different types of liquid biopsy features into a single model. Moreover, 4 additional papers identified through reference lists and manual search were added because they fulfilled the selection criteria. Only 25 articles were selected. **Table S1** presents the characteristics of these studies. Among them, 5 were prospective studies, and 18 were retrospective. The outcomes of these studies were mainly early cancer detection (19 articles), including multi-cancer early detection (MCED). The methods used for model development varied between the studies, and only 6 performed external validation in a completely independent dataset or study (temporal, technical, geographical, or domain). The most recent studies are the ones implementing this in their research. Neither calibration nor clinical utility was assessed in any of the studies.

This search was intended to provide a representative sample rather than a systematic or exhaustive review. It was limited to a single database (PubMed) and predefined search terms, so some relevant studies, particularly those using alternative terminology for multimodal, multiomics, or multi-analyte integration, may not have been captured. No PRISMA flow diagram was generated, and formal inclusion/exclusion criteria beyond those described above were not applied.

**Table S1**. Characteristics of the articles identified in the literature search

| Paper | Objective | Cancer type | Study design | Covariates | Sample size/ Population | Body fluid | Liquid biopsy features | Integration methods | Model evaluation metric | Validation strategy |
|---|---|---|---|---|---|---|---|---|---|---|
| **Chabon et al., 2020** [1] | Early detection | Lung cancer | Retrospective case-control study | Gender, age | 254 patients and controls for model development and internal validation | Blood | cfDNA: mutations, CNV, methylation | Ensemble classifier using 5NN, 3NN, naive Bayes, LR, and DT | AUC, accuracy | External validation in an independent prospective cohort (temporal) |
| **Gerratana et al., 2021** [2] | Metastatic sites identification | Breast cancer | Retrospective cohort study | Cancer stage, subtypes | 88 patients with metastatic breast cancer | Blood | CTCs and ctDNA: SNVs, INDELs, gene fusions/rearrangements, CNVs | Multivariate logistic regression, RF | Accuracy, mean per-class error | Internal validation: train-test split |
| **Li et al., 2021** [3] | Early detection | Liver cancer | Preclinical study | Tumor size | 16 HCC patients, 8 HBV carriers and 32 HC | Blood | cfDNA: methylation and sequencing | DL model | Accuracy and robustness | Internal validation: train-test split |
| **Ma et al., 2021** [4] | Early detection | Colorectal cancer | Retrospective case-control study | Gender, age | 390 CRC patients and 231 HC | Blood | cfDNA: fragmentomics, end motifs, CNV | Ensemble stacked model (using 5 ML models: GLM, GBM, RF, DL, XGBoost) | ROC curve, AUC, sensitivity, specificity | Internal validation: train-test split |
| **Hallermayr et al., 2022** [5] | Early detection and treatment response | Colorectal cancer | Retrospective case-control study | Cancer stage | 50 CRC patients and 61 HC | Blood | ctDNA: fragmentomics and CNV | ML classifier | ROC curve, AUC, sensitivity, specificity | Internal validation: train-test split |
| **Nakamura et al., 2022** [6] | Early detection | Pancreatic cancer | Retrospective cohort study | Cancer stage | 168 PDAC patients and 124 HC | Blood | Cell-free circulating miRNA, exosomal miRNA and CA19-9 | ML algorithms and quantitative real-time PCR assays | AUC | Internal validation: train-test split |
| **Wang et al., 2022** [7] | Early detection and TOO classification | Lung cancer | Prospective case-control study | Gender, age, nodule size, smoking status | 958 treatment naïve LUAD patients and 1318 HC | Blood | SMFs and CEA | ML models and DL | AUC, sensitivity, specificity | Internal validation: train-test split |
| **Eslami-S et al., 2023** [8] | Prognosis prediction | Lung cancer | Prospective cohort study | Treatment, tumor | 54 NSCLC patients | Blood | CTCs: presence/absence and EVs: PD-L1 | Multivariate Cox model | NA | No validation |

| | | | | | | | | | | |
|---|---|---|---|---|---|---|---|---|---|---|
| | | | | histological type | | | | | | |
| **Genco et al., 2023** [9] | Early detection | Pancreatic cancer | Preclinical study | NA | 47 patients | Cyst fluid and blood | cfDNA: mutations and proteins | KNN algorithm | Sensitivity, specificity | Internal validation: train-test split |
| **Haan et al., 2023** [10] | Early detection | Pancreatic cancer | Prospective case-control study | Age, gender, smoking status, stage, BMI, family history, diabetes | 4408 individuals (non-cancer, pancreatic cancer and precursor lesions) | Blood | cfDNA: methylation, fragmentation | Elastic Net | AUC, sensitivity | Internal validation: train-test split. Additional internal validation in other diseases from the same cohort |
| **Halner et al. 2023** [11] | MCED and early detection of lung cancer | Lung,breast, colorectal, oesophageal, liver, ovarian, pancreatic, gastric | Retrospective case-control study | Age, sex, ethnicity | MCED: 1005 cancer patients, 812 controls. Lung cancer: 61 lung cancer, 81 HC | Blood | Protein concentrations, cfDNA: DNA mutation 'omega' score | Data augmentation (KDE-based), feature selection (RF variable importance), multiple classifiers (RF, SVM, MLP, LR), classifier/hyperparameter optimization: Monte Carlo CV | ROC curve, AUC, sensitivity, specificity | Internal validation: hold-out test set (20%). Second external dataset to assess generalizability |
| **Kwon et al., 2023** [12] | Early detection | Lung cancer | Retrospective case-control study | NA | 92 lung cancer patients and 80 HC | Blood | cfDNA: percentage, CNV | Ensemble classifier | AUC | Internal validation: nested CV |
| **Nguyen et al., 2023** [13] | Early detection and TOO classification | Multi-cancer | Retrospective case-control study | Gender, age, cancer stage, tumor size | 738 non-metastatic patients (multi-cancer) and 1550 HC | Blood | cfDNA: methylation, CNA, fragments, end motifs | Ensemble stacking model using LR, and GCNN | AUC | Internal validation: train-test split |
| **Pham et al., 2023** [14] | Early detection and TOO classification | Breast cancer | Retrospective case-control study | NA | 239 breast cancer patients and 278 HC | Blood | cfDNA: methylation, CNA, end motifs | XGBoost | ROC curve, AUC | Internal validation: train-test split |
| **Wong et al., 2023** [15] | Prognosis prediction | Liver cancer | Retrospective case-control study | Clinicopathologic variables (not detailed) | 158 HCC patients and 33 HC | Blood | miRNA and proteins | RF | AUC, sensitivity, specificity | Internal validation: train-test split |

| | | | | | | | | | | |
|---|---|---|---|---|---|---|---|---|---|---|
| **Chen et al. 2024**[16] | Treatment response | Gastric cancer | Prospective longitudinal cohort study | Age, sex, smoking, Histology, Stage, Treatment | Training: 345 patients. External validation: 148 | Blood | CTC images and circulating proteins | Deep-learning-based dynamic-aware model | ROC curve, AUC | Internal validation: train-test-split |
| **Eledkawy et al., 2024** [17] | Early detection and TOO classification | Multi-cancer | Retrospective case-control study | Gender, age, race, histopatholog y | 1005 patients (multi-cancer) and 812 HC | Blood | cfDNA/ctDNA: mutations and protein concentration | XGBoost, LGMB | AUC, accuracy | (Partial) External validation with a publicly available dataset (geographical) |
| **Van Eijck et al., 2024** [18] | Treatment response | Pancreatic cancer | Prospective cohort study | Age, gender and disease stage | 58 PDAC patients | Blood | Plasma RNA and proteins | Stepwise logistic regression and BGLR | AUC, sensitivity, specificity | Internal validation: train-test split |
| **Zhao et al., 2024** [19] | Early detection | Pancreatic cancer | Retrospective cohort study | NA | 262 PDAC patients and 216 HC | Blood | cfDNA: methylation and CA19-9 | RF | ROC curve, AUC | Internal validation: train-test split |
| **Abraham et al. 2025**[20] | MCED, Genomic Probability Score (GPS) for diagnostic pathways, MRD and recurrence monitoring | Multi-cancer (solid tumors) | Observational, retrospective validation using multiple independent cohorts | Clinical stage, age, tumor fraction, survival outcomes, TOO, CHIP subtraction results, demographics | MCED training: 1,013 patients. Validation: 2,675 GPS training: 660 stage I/II + 506 normals (5-fold CV) MRD training: 3,439. Validation 86 (MRD) and 101 (monitoring) | Blood | Tissue: CNVs/ aneuploidy, SNVs/INDELs, fusions. LB: fragment length/dynamics, fragment diversity, nucleosome and TF footprint, end-motif, transcriptomics | XGBoost across nine feature pillars | Sensitivity, specificity, accuracy | External validation in multiple independent cohorts (geographical) for the different objectives |
| **Peng et al. 2025**[21] | Early detection | Renall cell carcinoma (RCC) | Retrospective case-control study | NA | Train: 280 participants (142 RCC, 138 HC). Validation: 161 (81 RCC, 81 HC). External validation: 230 (90 RCC, 140 HC) | Blood | ctDNA features: CNV, fragment size distribution and nucleosome footprint | Stacked ensemble model of individual base models (GLM, RF, NN and XGBoost) | ROC curve, AUC, sensitivity, specificity | External validation in an independent cohort (geographical) |
| **Tokareva et al. 2025**[22] | Early detection | Ovarian cancer | Retrospective case-control study | NA | 229 participants (103 serous ovarian cancer, 107 benign cases, 19 HC) | Blood | Lipidomics and metabolomics | CNN model based on Mann-Whitney-selected features, XGBoost + SVM-RFE | ROC curve, AUC, accuracy, | Internal validation |

| | | | | | | | | | | |
|---|---|---|---|---|---|---|---|---|---|---|
| | | | | | | | | | sensitivity, specificity | |
| **Van et al. 2025**[23] | Early detection | Breast cancer | Retrospective case-control study (training) and prospective cohort (validation) | NA | Train: 515 female participants (273 breast cancer, 108 BBD, 134 HC). External validation: 119 females (76 breast cancer, 43 BBD) | Blood | cfDNA: mutations, CNVs, methylation | Bootstrapping (model stability assessment). SelektKBest (feature selection). Five machine learning algorithms: SVM, DT, RF and XGBoost | ROC curve, AUC | External validation in an independent prospective cohort (temporal) |
| **Wang et al. 2025**[24] | Early detection | Oesophageal cancer | Retrospective case-control study | NA | 35 oesophageal cancer patients, 110 HC | Blood | cfDNA: methylation, fragmentomics | LASSO, RF | ROC curve, AUC, sensitivity, specificity, MCC, F1-score, balanced accuracy | Internal validation: train-test split |
| **Wang et al. 2026**[25] | Prognosis prediction | Non-small cell lung cancer | Retrospective cohort study | Age, sex, smoking, histology, stage, treatment | Training: 345 patients. External validation: 148 | Blood | VAF, ctDNA abundance, MRD, ctDNA mutations | NN, DT, random undersampling boosting, naïve Bayes, SVM and KNN | ROC curve, AUC, R1-score, accuracy, precision and recall | External validation in an independent prospective cohort (geographical) |

*AUC, Area Under the Curve; BBD, benign breast disease; BGLR, Bayesian Generalized Linear Regression; BMI, body mass index; CA19-9, carbohydrate antigen 19-9; CEA, carcinoembryonic antigen; cfDNA, cell-free DNA; CNVs, copy number variations; CRC, colorectal cancer; CTCs, circulating tumor cells; ctDNA, circulating tumor DNA; DL, Deep learning; DT, decision tree; EVs, extracellular vesicles; GBM, Gradient Boosting Machine; GCNN, Graph Convolutional Neural Network; GLM, Generalized Linear Model; GPS, genomic probability score; HBV, hepatitis B virus; HC, Healthy controls; HCC, Hepatocellular carcinoma; INDELs, insertions/deletions; KDE, kernel density estimation; KNN, K-Nearest Neighbors; LASSO, least absolute shrinkage and selection operator; LGBM: light gradient boosting machine; LR, Logistic regression; LUAD, lung adenocarcinoma; MCC, Matthews correlation coefficient; MCED, multi-cancer early detection; miRNA, micro-RNA; ML, machine learning; MLP, multi-layer perceptron; MRD, minimal residual disease; NN, neural network; NSCLC, non-small cell lung cancer; PDAC, pancreatic ductal adenocarcinoma; RF, Random forest; RCC; renal cell carcinoma; RFE, recursive feature elimination; ROC, Receiver Operating Characteristic; SMFs, serum metabolic fingerprints; SNV, single nucleotide variants; SVM, support vector machine; TF, transcription factor; TOO, tissue of origin; VAF, variant allele frequency; XGBoost, extreme gradient boosting.*

**Table S2**: Study designs for different LB study objectives: proposed designs, advantages, and limitations.

| Objective | | Proposed study design(s) | Pros | Cons | Examples from the selected papers (Table 1) |
|---|---|---|---|---|---|
| ***Screening*** | Discovery | Case-control | Utilizes existing data. Fast and efficient | No information on temporality. Potential for missing or incomplete data; prone to confounding, bias and batch effect | |
| | Validation | Prospective population cohort | Accounts for temporality | Less efficient and time-consuming | |
| | | Nested case-control | Accounts for temporality. More efficient than a prospective cohort | Difficult to obtain if the disease has low prevalence | |
| ***(Early) Detection of a symptomatic disease*** | Discovery | Case-control | Utilizes existing data. Fast and efficient. Enables biomarker discovery | Does not reflect the population at risk (healthy controls are often used) | Chabon et al., 2020 [1]<br>Ma et al., 2021 [4]<br>Hallermayr et al., 2022 [5]<br>Kwon et al., 2023[12]<br>Wong et al., 2023[15] |
| | Validation | Case series (in a population with suspicion of disease) | Can help validate the biomarker under realistic patient conditions | Requires a prolonged case accrual period | |
| ***Detection of tumor origin*** | Discovery/validation | Case series (in patients with diverse cancer types) | Can help identify and validate biomarkers that identify the tumor origin | Requires a prolonged case accrual period. | Nguyen et al., 2023[13]<br>Pham et al., 2023 [14]<br>Eledkawy et al. 2024[17] |
| ***Prognosis*** | Discovery/validation | Case series (in patients with cancer) | Can identify prognostic factors and establish temporality | May require extended follow-up, increasing the risk of loss to follow-up | |
| ***Treatment response*** | Discovery | Case series | Enables exploration of treatment response in real-world settings | Prone to bias; treatment details may be inconsistent or incomplete | |
| | Validation | Randomised clinical trials | Gold standard for assessing treatment efficacy; minimizes confounding | Costly and resource-intensive; may not always be feasible due to logistical, ethical or recruitment constraints | |
| ***MRD detection*** | Discovery | Case series | Enables exploration of MRD in real-world clinical settings; useful for early biomarker identification | Prone to selection bias and limited ability to assess prognostic performance or clinical utility | |

| | | | | |
|---|---|---|---|---|
| | Validation | Randomised clinical trials | Allows comparison of MRD-guided interventions versus standard-of-care to evaluate their efficacy. | Costly and resource-intensive; may face ethical and logistical challenges; not always feasible in rare populations |
| ***Uncovering resistance mechanisms*** | Discovery/validation | Case series (in patients with cancer and under treatment) | Enables exploration of resistance mechanisms in real-world settings | Prone to bias; data might be inconsistent or incomplete |
| ***Therapeutic targets identification*** | Discovery | Case series (in patients with cancer) | Can help identify therapeutic targets for posterior validation | More time-consuming to conduct |
| | Validation | Randomised clinical trials | Gold standard for assessing treatment efficacy; minimizes confounding | Costly and time-intensive; ethical considerations if withholding therapy from control groups |

**Table S3**: Liquid Biopsy features characteristics

*Bp, basepairs; CNV, Copy number variation; CPM: counts per million; ctDNA, circulating tumor DNA; FPKM, fragments per kilobase of transcript per million*

| | Feature | Type of variable | Measurement Range | Data Quantity | Distribution | Reference |
|---|---|---|---|---|---|---|
| QUANTIFICATION | **Concentration Cell/Particle Count Presence/Absence** | Quantitative (cell count), presence/absence, concentration | 0 to several hundred cells per mL | Low (a few to a few hundred cells) | Skewed with potential outliers | Lim et al., 2019[26] |
| *PROFILING* | **Point Mutations** | Mutational frequency, VAF | Variant allele frequency (VAF): 0.1% to >50% | Tens to hundreds of mutations | Continuous, variable range | Newman et al., 2016[27] |
| | **Copy Number Variations (CNVs)** | Copy number changes | $Log_2$ ratios or absolute copy numbers | Dozens to hundreds of CNVs | Continuous, variable range | Heitzer, E., 2015[28] |
| | **Transcriptomics** | Gene expression levels | CPM, TPM or FPKM | High (thousands of genes/transcripts) | Continuous, expression level variability | Ignatiadis et al., 2021[29] |
| | **Proteomics** (including secretomics) | Protein expression levels | Relative abundance or concentration | High (hundreds to thousands of proteins) | Continuous, variable range | Zhang et al., 201[30] |
| | **Metabolomics** | Metabolite concentrations | Nanomolar to millimolar concentrations | Moderate to high (hundreds to thousands of metabolites) | Continuous, nanomolar to millimolar | Wishart, D., 2016[31] |
| | **Fragmentomics** | Fragment size, end motifs, nucleosome occupancy | Fragment length < 200 bp for ctDNA, end motifs: sequence-specific | Low abundance of ctDNA, from ≥ 5–10% in late stage to ≤ 0.01–0.1% in early-stage cancers | Continuous (fragment lengths), categorical (end motifs) | Havell et al., 2022 [32] Lo et al. 2021[33] |
| | **Epigenomics** | DNA methylation (M), histone modifications, nucleosome positioning (N) | M: 0 to 100% (or 0-1) N: Varies by signal (e.g., normalized coverage depth, peak scores); indirectly reflects nucleosome occupancy | M: High (hundred thousands of CpG sites/histone marks). Twist panel (targeted) has less than whole genome methylation/ N: Moderate to High (depends on sequencing depth) | M: Continuous, 0 to fully methylated 1 N: Continuous (coverage signal, position-based metrics like NFR scores) | Guo et al., 2017[34] Vanderstichele et al. 2022 [35] |

*mapped reads; M, methylation; N, nucleosome positioning; NFR, nucleosome-free region; TPM, transcripts per million; VAF, variant allele frequency.*

**Table S4**: Summary of batch effect correction methods: characteristics, advantages, and limitations for application in liquid biopsy.

| Method | Input data | Description | Pros | Cons | Package / Reference |
|---|---|---|---|---|---|
| **Batch mean centering** | $log_2$-transformed expressions | Center data within each batch. Univariate method. | -Simple and computationally efficient<br>-Requires minimal assumptions | -Non-optimal for non-Gaussian data<br>-Limited correction scope; does not account for complex batch effects | Base R |
| **ComBat /ComBat-seq** | ComBat: Expressions of proteomics and metabolomics.<br>ComBat-seq: transcriptomics | ComBat: assumes a Gaussian distribution of the data<br>ComBat-seq: negative binomial regression to model batch effect, mapping the data to an expected batch-free distribution. | -Effective for Gaussian or count-based data<br>-Widely used and well-documented | -Performance depends on data distribution assumptions<br>-Sensitive to outliers and poor if assumptions are violated | Sva<br>https://github.com/zhangyuqing/ComBat-seq<br>Zhang et al. 2020[36] |
| **Harmony** | Originally developed for single-cell RNA-seq data, but adaptable to other omics. Starts with a low-dimensional embedding of cells (e.g., Principal Component Analysis). | Clusters cells based on initial embedding and adjusts the embedding iteratively moving cells towards a consensus position with cells from other batches that belong to the same cluster, maintaining biological signals, and removing batch-specific variation. | -Maintains biological signal integrity<br>-Effective for complex multi-batch datasets | -Computationally intensive.<br>-Requires appropriate low-dimensional embeddings as input | Harmony<br>https://github.com/immunogenomics/harmony<br>Korsunsky et al. 2019[37] |
| **Surrogate variable analysis** | Various types of expression data. Suitable for high-dimensional data and unknown batch origin. | Constructs surrogate variables (covariates from high-dimensional data) which are then used to adjust for unknown, unmodeled, or latent noise sources. | -Handles unknown and latent batch effects effectively<br>-Flexible for high-dimensional data | -Requires sufficient sample sizes for surrogate variable estimation<br>-May introduce additional variability if misapplied | SVA<br>https://bioconductor.org/packages/release/bioc/html/sva.html |
| **Remove unwanted variation** | Various types of expression data. Suitable for high-dimensional data. | Identifies and removes unwanted variation by using control genes or negative control samples. | -Effective for high-dimensional data<br>-Allows explicit control through reference or control genes/samples | -Relies on accurate identification of control genes/samples<br>-May struggle with unknown batch effects | Ruv<br>https://cran.r-project.org/web/packages/ruv/index.html |
| **Ratio-based scaling** | Various types of expression data. Used in proteomics by comparing the relative abundance of proteins across different batches. | It is obtained by subtracting $log_2$ transformed expression profiles of a feature by the mean of the $log_2$ transformed expression profile of the reference | -Simple and easy to implement<br>-Effective for adjusting relative differences in proteomics data | -May not fully correct for complex batch effects<br>-Relies on stable and representative reference features | Base R |

**Table S5**: Summary of model performance metrics: purpose, equation, advantages, and disadvantages.

| Metric | Purpose | Equation | Pros | Cons |
|---|---|---|---|---|
| **Accuracy** | Classification | $\frac{TP + TN}{TP + FP + TN + FN}$ | -Simple and intuitive<br>-Works well on balanced datasets | -Misleading on imbalanced datasets.<br>-Does not differentiate between FP and FN |
| **Error rate** | Classification | $\frac{FP + FN}{TP + FP + TN + FN}$<br>$1 - Accuracy$ | -Simple and intuitive<br>-Works well on balanced datasets | -Misleading on imbalanced datasets<br>-Does not differentiate between FP and FN |
| **Sensitivity/ recall/TPR** | Classification | $\frac{TP}{TP + FN}$ | -Emphasizes true positive detection<br>-Useful for rare-event detection | -Ignores FP<br>-Can give a false sense of performance if precision is low |
| **Specificity/ TNR** | Classification | $\frac{TN}{TN + FP}$ | -Emphasizes true negative detection<br>-Useful when negatives are important to classify | -Ignores FN<br>-Can give a false sense of performance if sensitivity is low |
| **Precision/ PPV** | Classification | $\frac{TP}{TP + FP}$ | -Focuses on relevance of positive predictions<br>-Useful when FP are costly | -Ignores FN<br>-Can give misleading performance if sensitivity is low<br>-Depends on disease prevalence |
| **NPV** | Classification | $\frac{TN}{TN + FN}$ | -Focuses on relevance of negative predictions<br>-Useful when FN are costly | -Ignores FP<br>-Can give misleading performance if specificity is low<br>-Depends on disease prevalence |
| **FPR** | Classification | $\frac{FP}{FP + TN}$<br>$1 - Specificity$ | -Helps understand false alarm rates<br>-Complements specificity | -Alone, it does nott provide information about positive class detection |
| **FNR** | Classification | $\frac{FN}{FN + TP}$<br>$1 - Sensitivity$ | -Helps understand missed detection rates<br>-Complements sensitivity | -Alone, it does not provide information about negative class detection |
| **F1-score** | Classification | $2 \times \frac{Precision \times Recall}{Precision + Recall}$ | -Balances precision and recall<br>-Suitable for imbalanced datasets | -Doesn't differentiate between the cost of FP and FN |
| **ROC curve** | Classification | *Sensitivity vs. 1 - Specificity* | -Summarizes classification performance across threshold<br>-Useful for balanced datasets | -Misleading on imbalanced datasets<br>-Focuses on ranking predictions rather than absolute performance |

| | | | | |
|---|---|---|---|---|
| **AUC-ROC** | Classification | $AUC = \int_0^1 TPR(FPR)d(FPR)$ | -Threshold independent<br>-Provides a single number summary | -Misleading with class imbalance<br>-Treats FPR uniformly |
| **PR curve** | Classification | *Precision vs. Recall* | -Better for imbalanced datasets<br>-Highlights trade-off between precision and recall | -Ignores true negatives<br>-Can be harder to interpret compared to ROC |
| **AUC_PR** | Classification | $AUC = \int_0^1 Precision(Recall)d(Recall)$ | -Very informative for highly imbalanced datasets<br>-Focuses on positive-class performance | -Harder to interpret than ROC-AUC<br>-Baseline depends on class prevalence |
| **Youden's Index** | Classification | $Sensitiviy + Specificity - 1$ | -Combines sensitivity and specificity<br>-Helps identify the optimal threshold | -Oversimplifies by equally weighting sensitivity and specificity |
| **G-mean** | Classification | $G - mean = \sqrt{Sensitivity * Specificity}$ | -Useful for imbalanced datasets, not dominated by the majority class<br>-Encourages classifiers that perform well on both minority and majority classes | - Does not differentiate between the cost of FP and FN |
| **MCC** | Classification | $\frac{(TP \times TN) - (FP \times FN)}{\sqrt{(TP + FP)(TP + FN)(TN + FP)(TN + FN)}}$ | -Balanced metric considering all confusion matrix elements<br>-Works well for imbalanced datasets | -Computationally intensive<br>-Hard to interpret compared to simpler metrics like accuracy |
| **$LR^+$** | Classification | $\frac{Sensitivity}{1 - Specificity}$ | -Quantifies how much a positive result increases disease likelihood | -Requires both sensitivity and specificity<br>-May be challenging to interpret directly |
| **$LR^-$** | Classification | $\frac{1 - Sensitivity}{Specificity}$ | -Quantifies how much a negative result decreases disease likelihood | -Requires both sensitivity and specificity<br>-May be challenging to interpret directly |
| **DOR** | Classification | $\frac{LR^+}{LR^-}$<br>$\frac{TP \times TN}{FP \times FN}$ | -Combines sensitivity and specificity into a single number | -Does not provide information about specific error types |
| **Micro-AUC** | Classification | AUC−ROC obtained by aggregating all class predictions and labels across classes | -Aggregates all predictions<br>-Works well for overall performance | -Dominated by majority classes, making it less informative for imbalanced datasets |
| **Macro-AUC** | Classification | $\frac{1}{k}\sum_{i=1}^{k} AUC_i$ | -Treats all classes equally<br>-Suitable for comparing performance across classes | -Ignores class imbalance<br>-May overemphasize minority classes with poor performance |

| | | | | |
|---|---|---|---|---|
| **MAE** | Regression | $\frac{1}{n}\sum_{i=1}^{n}\lvert y_i - ypred_i\rvert$ | -Easy to interpret; errors are in the same unit as the original data<br>-Robust to outliers compared to MSE and RMSE<br>-Provides a straightforward average error magnitude | -Does not penalize large errors more than small errors (lacks quadratic emphasis)<br>-May not be suitable when large errors need greater attention |
| **MSE** | Regression | $\frac{1}{n}\sum_{i=1}^{n}(y_i - ypred_i)^2$ | -Penalizes large errors more than small ones, making it useful for capturing significant deviations<br>-Differentiable, making it suitable for optimization in machine learning | -Not in the same units as the original data (squared scale)<br>-Sensitive to outliers, as large errors are squared |
| **RMSE** | Regression | $\sqrt{\frac{1}{n}\sum_{i=1}^{n}(y_i - ypred_i)^2}$ | -In the same units as the original data, making it easier to interpret than MSE<br>-Penalizes large errors like MSE, which is useful when outliers are important | -Still sensitive to outliers due to squaring errors<br>-More computationally expensive because of the square root |
| **$R^2$** | Regression | $1 - \frac{\sum_{i=1}^{n}(y_i - ypred_i)^2}{\sum_{i=1}^{n}(y_i - \overline{y})^2}$ | -Indicates how well the model explains the variance in the target variable<br>-Normalized between 0 and 1 for easy interpretation | -Can give misleading results if the model is overfitted or if the data is non-linear<br>-Not intuitive for understanding absolute error magnitudes |
| **Adjusted $R^2$** | Regression | $1 - \left(\frac{(1-R^2)(n-1)}{n-k-1}\right)$ | -Penalizes overfitting by accounting for the number of predictors<br>-Provides a more realistic measure of model performance for multiple predictors | -Can still overestimate performance if irrelevant predictors are included<br>-Slightly more complex to calculate and interpret compared to $R^2$ |

AUC, area under the curve; DOR, diagnostic odds ratio; $LR^+$, positive diagnostic likelihood ratio; $LR^-$, negative diagnostic likelihood ratio; FN, false negative; FNR, false negative rate; FP, false positive; FPR, false positive rate; k, number of classes; MAE, mean absolute error; MCC, Matthews Correlation Coefficient; MSE, mean square error; n, number of samples; NPV, negative predictive value; PPV, positive predictive value; PR, precision-recall; $R^2$, R squared; RMSE, root mean squared error; ROC, receiver operation characteristic; TN, true negative; TNR, true negative rate; TP, true positive; TPR, true positive rate

**Table S6**: Summary of the model-agnostic methods to interpret ML models.

| | Method | Model Explanation | Pros | Cons |
|---|---|---|---|---|
| *Global methods* | Partial dependence plot | Shows the marginal effect of 1 or 2 features on the predicted outcome | -Easy to interpret<br>-Useful for assessing feature importance | -Assumes independence between features, which may not hold in practice<br>-Limited to 1-2 features |
| | Accumulated local effects plot | Describes how a feature affects the average prediction | -Accounts for feature dependencies<br>-Computationally efficient compared to PDP | -Still limited to feature-specific effects<br>-May miss nuanced interactions |
| | Feature interactions | H-Statistic. Measures the interaction between two variables or between one variable and the rest. It has meaningful interpretation and works with all kinds of interactions. Computationally expensive | -Quantifies interactions effectively<br>-Broad applicability across interaction types | -High computational cost for large datasets<br>-Difficult to interpret with complex interactions |
| | Functional decomposition | Deconstructs the high-dimensional function as a sum of individual effects and interaction effects | -Provides a detailed breakdown of feature contributions<br>-Captures complex interactions | -Challenging to implement<br>-Interpretation becomes harder with increasing dimensions |
| | Permutation feature importance | Analyzes the increase in prediction error after permuting the feature's values | -Provides clear feature importance rankings | -Sensitive to correlated features<br>-Can underestimate the importance of features involved in complex interactions |
| | Global surrogate models | Trains interpretable models to approximate the predictions of a black-box model | -Intuitive approach to explainability<br>- Works with any black-box model | -May lose fidelity when approximating complex models<br>-Choice of surrogate impacts interpretability |
| | Prototypes and criticisms | Maximum Mean Discrepancy. It finds prototypes (well-representative data instances) and criticisms (data instances not well-represented by prototypes) | -Helps identify representative and outlier instances<br>-Offers insights into data structure | -Computationally expensive for large datasets<br>-Prototype selection can be subjective |

| | | | | |
|---|---|---|---|---|
| *Local Methods* | Individual conditional expectation | Visualizes how the prediction of a ML model changes for an individual instance as a specific feature varies | -Provides instance-level insights<br>-Highlights heterogeneous effects across instances | -Limited to one feature at a time<br>-Can be visually complex for high-dimensional data |
| | Local interpretable model-agnostic explanations (LIME) | Interpretable models are used to explain individual predictions made by a black-box model | -Simple, intuitive, and local interpretability<br>-Works for any model type | -Sensitive to sampling strategy<br>-May not generalize well to other instances |
| | Counterfactual explanations | Identifies how to minimally alter an individual instance's features to change its prediction to a desired outcome, providing insight into the model's decision boundary | -Useful for actionable insights<br>-Provides a clear sense of decision boundaries | -Computationally expensive for high-dimensional data<br>-May yield unrealistic or non-actionable suggestions |
| | Scoped rules (anchors) | Generates if-then rules (anchors), highlighting conditions under which a model consistently makes the same prediction | -Produces interpretable, human-readable rules<br>-Focuses on high-confidence decision regions | -Rule generation may be infeasible for highly complex models<br>-Rules may oversimplify decision logic |
| | Shapley values | Quantify each feature's contribution to a prediction by considering all possible feature combinations, offering a fair, cooperative game theory-based measure of feature importance | -Fair and theoretically sound (game-theoretic)<br>-Handles correlated features better than simpler importance measures | -Computationally expensive, especially for high-dimensional models |
| | SHAP | Combines Shapley values with local explanations, offering a fair measure of feature importance by attributing each feature's contribution to individual predictions | -Balances local and global interpretability<br>-Robust for understanding complex models | -Computational cost can be high for large datasets or models<br>-Interpretation may be challenging for non-experts |

LIME, local interpretable model-agnostic explanations; ML, machine learning; PDP, partial dependence plot; SHAP, Shapley additive explanations.

### References


1. Chabon JJ, Hamilton EG, Kurtz DM, et al. Integrating genomic features for non-invasive early lung cancer detection. *Nature*. 2020;580(7802):245-251. doi:10.1038/s41586-020-2140-0
2. Gerratana L, Davis AA, Polano M, et al. Understanding the organ tropism of metastatic breast cancer through the combination of liquid biopsy tools. *Eur J Cancer*. 2021;143:147-157. doi:10.1016/j.ejca.2020.11.005
3. Li J, Zhang X, Zhang W, et al. DISMIR: Deep learning-based noninvasive cancer detection by integrating DNA sequence and methylation information of individual cell-free DNA reads. *Brief Bioinform*. 2021;22(6). doi:10.1093/BIB/BBAB250
4. Ma X, Chen Y, Tang W, et al. Multi-dimensional fragmentomic assay for ultrasensitive early detection of colorectal advanced adenoma and adenocarcinoma. *J Hematol Oncol*. 2021;14(1). doi:10.1186/s13045-021-01189-w
5. Hallermayr A, Wohlfrom T, Steinke-Lange V, et al. Somatic copy number alteration and fragmentation analysis in circulating tumor DNA for cancer screening and treatment monitoring in colorectal cancer patients. *J Hematol Oncol*. 2022;15(1). doi:10.1186/s13045-022-01342-z
6. Nakamura K, Zhu Z, Roy S, et al. An exosome-based transcriptomic signature for noninvasive, early detection of patients with pancreatic ductal adenocarcinoma: A multicenter cohort study. *Gastroenterology*. 2022;163(5):1252-1266.e2. doi:10.1053/j.gastro.2022.06.090
7. Wang L, Zhang M, Pan X, et al. Integrative serum metabolic fingerprints based multi-modal platforms for lung adenocarcinoma early detection and pulmonary nodule classification. *Adv Sci (Weinh)*. 2022;9(34):2203786. doi:10.1002/advs.202203786

8. Eslami-S Z, Cortés-Hernández LE, Sinoquet L, et al. Circulating tumour cells and PD-L1-positive small extracellular vesicles: the liquid biopsy combination for prognostic information in patients with metastatic non-small cell lung cancer. *Br J Cancer*. 2023;130(1):63-72. doi:10.1038/s41416-023-02491-9
9. Genco E, Modena F, Sarcina L, et al. A single-molecule bioelectronic portable array for early diagnosis of pancreatic cancer precursors. *Adv Mater*. 2023;35(42):2304102. doi:10.1002/adma.202304102
10. Haan D, Bergamaschi A, Friedl V, et al. Epigenomic blood-based early detection of pancreatic cancer employing cell-Free DNA. *Clinical Gastroenterology and Hepatology*. 2023;21(7):1802-1809.e6. doi:10.1016/j.cgh.2023.03.016
11. Halner A, Hankey L, Liang Z, et al. DEcancer: Machine learning framework tailored to liquid biopsy based cancer detection and biomarker signature selection. *iScience*. 2023;26(5). doi:10.1016/j.isci.2023.106610
12. Kwon HJ, Park UH, Goh CJ, et al. Enhancing lung cancer classification through integration of liquid biopsy multi-omics data with machine learning techniques. *Cancers (Basel)*. 2023;15(18). doi:10.3390/cancers15184556
13. Nguyen VTC, Nguyen TH, Doan NNT, et al. Multimodal analysis of methylomics and fragmentomics in plasma cell-free DNA for multi-cancer early detection and localization. 2023;12. doi:10.7554/elife.89083
14. Pham TMQ, Phan TH, Jasmine TX, et al. Multimodal analysis of genome-wide methylation, copy number aberrations, and end motif signatures enhances detection of early-stage breast cancer. *Front Oncol*. 2023;13. doi:10.3389/fonc.2023.1127086
15. Wong VCL, Wong MI, Lee VHF, Man K, Ng KTP, Cheung TT. Prognostic MicroRNA fingerprints predict recurrence of early-stage hepatocellular carcinoma following hepatectomy. *J Cancer*. 2023;14(3):480-489. doi:10.7150/jca.79593

16. Chen Z, Zhao J, Li Y, et al. Predicting response to patients with gastric cancer via a dynamic-aware model with longitudinal liquid biopsy data. *Gastric Cancer*. 2025;28(5):886-898. doi:10.1007/s10120-025-01628-4
17. Eledkawy A, Hamza T, El-Metwally S. Precision cancer classification using liquid biopsy and advanced machine learning techniques. *Sci Rep*. 2024;14(1). doi:10.1038/s41598-024-56419-1
18. van Eijck CWF, Sabroso-Lasa S, Strijk GJ, et al. A liquid biomarker signature of inflammatory proteins accurately predicts early pancreatic cancer progression during FOLFIRINOX chemotherapy. *Neoplasia*. 2024;49. doi:10.1016/j.neo.2024.100975
19. Zhao G, Jiang R, Shi Y, et al. Circulating cell-free DNA methylation-based multi-omics analysis allows early diagnosis of pancreatic ductal adenocarcinoma. *Mol Oncol*. Published online 2024. doi:10.1002/1878-0261.13643
20. Abraham J, Domenyuk V, Perdigones N, et al. Validation of an AI-enabled exome/transcriptome liquid biopsy platform for early detection, MRD, disease monitoring, and therapy selection for solid tumors. *Scientific Reports 2025 15:1*. 2025;15(1):21173-. doi:10.1038/s41598-025-08986-0
21. Peng YL, Yu B, Huang TX, et al. Early detection of renal cell carcinoma: a novel cell-free DNA fragmentomics-based liquid biopsy assay. *ESMO Open*. 2025;10(7). doi:10.1016/j.esmoop.2025.105323
22. Tokareva A, Iurova M, Starodubtseva N, et al. Machine learning framework for ovarian cancer diagnostics using plasma lipidomics and metabolomics. *International Journal of Molecular Sciences 2025, Vol 26,*. 2025;26(14). doi:10.3390/ijms26146630
23. Van TTV, Tran TH, Nguyen THH, et al. Multimodal analysis of cell-free DNA enhances differentiation of early-stage breast cancer from benign lesions and healthy individuals. *BMC Biol*. 2025;23(1). doi:10.1186/s12915-025-02371-z

24. Wang J, Zhang W, Li H, Wang Y, Qi J, Nie J. Cell-free DNA methylation and fragmentomics-based liquid biopsy for accurate esophageal cancer detection. *BMC Cancer*. 2025;25(1). doi:10.1186/s12885-025-15150-4
25. Wang Y, Xiang YB, Chen XW, et al. PRIME: an interpretable artificial intelligence model based on liquid biopsy improves prediction of progression risk in non-small cell lung cancer. *Mil Med Res*. 2025;12(1). doi:10.1186/s40779-025-00679-z
26. Lim S Bin, Di Lee W, Vasudevan J, Lim WT, Lim CT. Liquid biopsy: one cell at a time. *npj Precision Oncology 2019 3:1*. 2019;3(1):1-9. doi:10.1038/s41698-019-0095-0
27. Newman AM, Lovejoy AF, Klass DM, et al. Integrated digital error suppression for improved detection of circulating tumor DNA. *Nat Biotechnol*. 2016;34(5):547-555. doi:10.1038/nbt.3520,
28. Heitzer E, Ulz P, Geigl JB. Circulating tumor DNA as a liquid biopsy for cancer. *Clin Chem*. 2015;61(1):112-123. doi:10.1373/clinchem.2014.222679,
29. Ignatiadis M, Sledge GW, Jeffrey SS. Liquid biopsy enters the clinic — implementation issues and future challenges. *Nature Reviews Clinical Oncology 2021 18:5*. 2021;18(5):297-312. doi:10.1038/s41571-020-00457-x
30. Zhang B, Wang J, Wang X, et al. Proteogenomic characterization of human colon and rectal cancer. *Nature 2014 513:7518*. 2014;513(7518):382-387. doi:10.1038/nature13438
31. Wishart DS. Emerging applications of metabolomics in drug discovery and precision medicine. *Nature Reviews Drug Discovery 2016 15:7*. 2016;15(7):473-484. doi:10.1038/nrd.2016.32
32. Markus H, Chandrananda D, Moore E, et al. Refined characterization of circulating tumor DNA through biological feature integration. *Sci Rep*. 2022;12(1):1-11. doi:10.1038/s41598-022-05606-z

33. Lo YMD, Han DSC, Jiang P, Chiu RWK. Epigenetics, fragmentomics, and topology of cell-free DNA in liquid biopsies. *Science (1979)*. 2021;372(6538). doi:10.1126/science.aaw3616,

34. Guo S, Diep D, Plongthongkum N, Fung HL, Zhang K, Zhang K. Identification of methylation haplotype blocks aids in deconvolution of heterogeneous tissue samples and tumor tissue-of-origin mapping from plasma DNA. *Nat Genet*. 2017;49(4):635-642. doi:10.1038/ng.3805

35. Vanderstichele A, Busschaert P, Landolfo C, et al. Nucleosome footprinting in plasma cell-free DNA for the pre-surgical diagnosis of ovarian cancer. *NPJ Genom Med*. 2022;7(1):1-9. doi:10.1038/s41525-022-00300-5

36. Zhang Y, Parmigiani G, Johnson WE. ComBat-seq: batch effect adjustment for RNA-seq count data. *NAR Genom Bioinform*. 2020;2(3). doi:10.1093/nargab/lqaa078

37. Korsunsky I, Millard N, Fan J, et al. Fast, sensitive and accurate integration of single-cell data with Harmony. *Nature Methods 2019 16:12*. 2019;16(12):1289-1296. doi:10.1038/s41592-019-0619-0
